# Molecular dynamics simulations illuminate the role of sequence context in the ELF3-PrD-based temperature sensing mechanism in plants

**DOI:** 10.1101/2023.03.15.532793

**Authors:** Richard J. Lindsay, Abhilash Sahoo, Rafael Giordano Viegas, Vitor B. P. Leite, Philip A. Wigge, Sonya M. Hanson

**Affiliations:** Center for Computational Biology, Flatiron Institute, New York, NY 10010, USA; Center for Computational Mathematics, Flatiron Institute, New York, NY 10010, USA; Department of Physics, São Paulo State University (UNESP), Institute of Biosciences, Humanities and Exact Sciences, São José do Rio Preto, SP, 15054-000, Brazil; Federal Institute of Education, Science and Technology of São Paulo (IFSP), Catanduva, São Paulo 15.808-305, Brazil; Leibniz-Institut für Gemüse-und Zierpflanzenbau, Großbeeren, Germany; Institute of Biochemistry and Biology, University of Potsdam, Potsdam, Germany

## Abstract

The evening complex (EC) is a tripartite DNA repressor and a core component of the circadian clock that provides a mechanism for temperature-responsive growth and development of many plants. ELF3, a component of the EC, is a disordered scaffolding protein that blocks transcription of growth genes at low temperature. At increased temperature EC DNA binding is disrupted and ELF3 is sequestered in a reversible nuclear condensate, allowing transcription and growth to proceed. The condensation is driven by a low complexity prion-like domain (PrD), and the sensitivity of the temperature response is modulated by the length of a variable polyQ tract, with a longer polyQ tract corresponding to enhanced condensate formation and hypocotyl growth at increased temperature. Here, a series of computational studies provides evidence that polyQ tracts promote formation of temperature-sensitive helices in flanking residues with potential impacts for EC stability under increasing temperature. REST2 simulations uncover a heat-induced population of condensation-prone conformations that results from the exposure of ‘sticky’ aromatic residues by temperature-responsive breaking of long-range contacts. Coarse-grained Martini simulations reveal both polyQ tract length and sequence context modulate the temperature dependence of cluster formation. Understanding the molecular mechanism underlying the ELF3-PrD temperature response in plants has implications for technologies including modular temperature-response elements for heat-responsive protein design and agricultural advances to enable optimization of crop yields and allow plants to thrive in increasingly inhospitable environments.

## Introduction

Temperature sensing mechanisms are critical to the survival of all life, but for plants, which are mostly stationary and yet flourish in a vast array of climates and seasonal variations, temperature sensors have a key, yet still poorly understood, role. While plants have diverse schemes to survive and thrive at a broad range of temperatures, such as vernalization, heat stress, and cold stress (Kerbler and Wigge, 2023), one mechanism that is more specific to plants is thermomorphogenesis, or when plants adjust their morphology in response to temperature (Quint et al., 2016).

Several thermosensory biomolecules have been identified, like the temperature-responsive uncoiling of chromatin, bending of DNA to increase gene transcription, and the melting of hairpin loops in RNA to enhance protein expression (Paik and Huq, 2019; Lin et al., 2020; Sengupta and Garrity, 2013). One specific component known to be involved in thermomorphogenesis is the Evening Complex (EC), whose function depends on protein-protein and protein-DNA interactions, and has been well-studied in *Arabidopsis thaliana*. The EC is conserved in land plants (Liu et al., 2001) and is a tripartite protein complex that provides a tunable temperature-sensing mechanism with implications for growth rate, flowering, and potentially senescence and circadian rhythm (Zagotta et al., 1992; Anwer et al., 2014; Nusinow et al., 2011; Thines and Harmon, 2010; Box et al., 2015; Jung et al., 2020). Understanding the molecular basis for the temperature sensing mechanism of the Evening Complex has implications on a warming planet where even a 1°C rise in temperature would affect crop yield (Zhao et al., 2017).

The EC is a transcription repressor with three known protein components – ELF3, ELF4, and LUX ARRYTHMO – where LUX is a transcription factor (Nusinow et al., 2011; Zhang et al., 2019), ELF4 is a small helical protein responsible for stabilizing the complex, and ELF3 is a large mostly disordered protein that sequesters into reversible nuclear condensates at high temperatures (Fig. 1A) resulting in the release of the rest of the repressor complex, allowing the expression of growth genes (Jung et al., 2020). While ELF3 is a 695 residue long intrinsically disordered protein (IDP) the C-terminal prion-like domain (PrD), first recognized by Jung et al. (2020)., is necessary and sufficient for condensate formation (Fig. 1C) and makes up a quarter of the ELF3 sequence (residues 492 to 664 out of 695 total residues) (Jung et al., 2020). Prion-like domains are identified by matching sequence patterns found initially in amyloid-forming prions, namely via an enrichment in polar amino acids like glutamine and asparagine (Lancaster et al., 2014), but this has proved a successful way of identifying low-complexity domains that are indicative of disordered protein regions (Romero et al., 2001). The PrD in ELF3 (hereafter referred to as the ELF3-PrD) is critically involved in the temperature sensitivity of the EC, as the ELF3 of the Mediterranean grass species *Brachypodium distachyon* has no PrD and does not undergo phase change or accelerated growth at higher temperatures (Jung et al., 2020).

**Figure 1.**
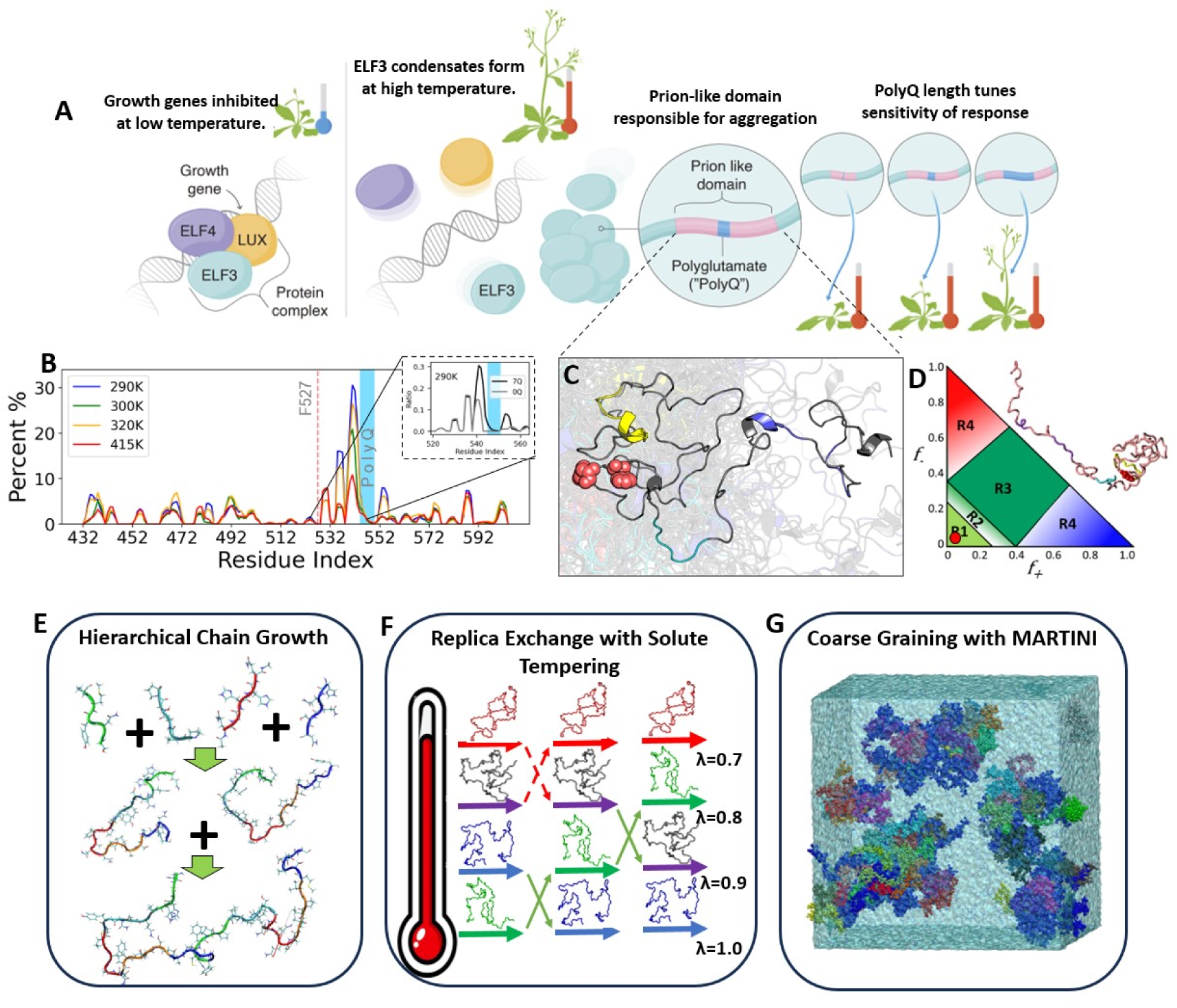
An overview of the Evening Complex (EC), key features of ELF3, and the computational approaches taken to elucidate the molecular mechanisms of temperature sensing of the ELF3-PrD. **A.** An illustration of the temperature-responsive mechanism of the evening complex. The left panel shows the intact EC repressing a growth gene target. Second to left illustrates the complex dissociating from DNA at high temperature. The next section describes the prion-like domain required for condensation and the poly-glutamine region responsible for tuning the sensitivity of the temperature response. The right-most cartoon illustrates the enhanced growth response characteristic of expanded poly-glutamine tracts. **B.** Helical propensity of hierarchical chain growth (HCG) ensembles by residue at four temperatures. The 7Q (wild-type) system is shown in the main panel with a region of interest of 7Q and 0Q shown in the inset. The light blue regions mark the locations of the poly-glutamine tracts and the yellow region highlights an aromatic helix of mechanistic importance. C. A structure of wild-type ELF3-PrD taken from a REST2 simulation. **D.** A charge fraction plot commonly used to classify IDPs based on sequence. The red dot indicates ELF3-PrD falls in region 1, classifying it as a weak polyampholite characterized by a globular or ’tadpole-like’ shape. **E.** An illustration of the hierarchical chain growth method used to produce ELF3-PrD ensembles at a variety of polyQ lengths and temperature conditions. **F.** A cartoon representation of the REST2 enhanced sampling simulation method. **G.** A snapshot from a coarse-grained Martini simulation used here to investigate temperature-dependent condensation propensity of ELF3-PrD variants.

A central feature of ELF3-PrD in *Arabidopsis thaliana* is the presence of a poly-glutamine (polyQ) tract of variable length found to differ in *Arabidopsis* populations by geographic location (Anwer et al., 2014). A similar polyQ tract expansion is responsible for the formation of pathogenic huntingtin protein aggregates in Huntington’s disease patients. However the tract lengths observed in Huntington’s disease are longer (>35 residues) than those of ELF3, which contains polyQ tracts of 0 to 29 glutamine residues. (Reiner et al., 2011). Experiments by Jung et al. found the length of this polyQ tract modulates the sensitivity of the temperature response, observing that longer polyQ enhances condensate formation and hypocotyl growth rate at high temperature (Jung et al., 2020). The mechanistic insights gained from studies of ELF3 condensate formation and temperature responsiveness could shed light on general principles governing polyQ-mediated interactions in native cellular contexts.

That evolution has co-opted biological condensates for temperature sensing in *Arabidopsis* aligns with the intrinsic temperature-dependence of condensate formation as well as the nature of IDPs as key sensors of cellular physicochemistry in general (Moses et al., 2023). However understanding the driving forces for IDP condensate formation at high (LCST) vs. low (UCST) temperatures is still just beginning to be understood. Here, condensate formation at higher temperatures, like ELF3-PrD, correspond to those with a lower critical solution temperature (LCST) and those that form condensates at lower temperatures have an upper critical solution temperature (UCST) – these abbreviations will be used from now on. The understanding of LCST vs. UCST behavior has improved recently, for example via sequence heuristics proposed in (Quiroz and Chilkoti, 2015; Ruff et al., 2018), but the understanding physicochemical drivers of native LCSTs is still a work in progress. Sequence heuristics in general are a useful way to understand IDP functions (Maristany et al., 2023), for example they can accurately identify classes of IDP by simply looking at fraction of positive and negative charges in the sequence (Fig. 1D) (Das and Pappu, 2013). However, understanding the interplay between the polyQ regions of ELF3-PrD and the other sequence heuristics available is non-trivial. Previous studies have shown that polyQ tracts often enhance helical propensity in adjacent regions (Totzeck et al., 2017), which is of particular interest because short regions of transient helicity called Short Linear Motifs (SLiMs) are often responsible for formation of intra- and inter-protein interactions necessary for biological function (Davey et al., 2012).

The dynamic nature of IDPs has historically complicated their structural characterization (Bari and Prakashchand, 2021; Schramm et al., 2019). Experimental approaches can provide information about ensemble averages, but elucidating information about specific states or clusters of states remains a challenge (Nag et al., 2022). Using computational approaches to bridge this gap and access conformational ensembles, the molecular characterization of condensates is becoming increasingly accessible. While atomistic molecular dynamics (MD) simulations offer the most precise access to these ensembles, the computational cost of sampling the full conformational space of an IDP, not to mention a condensate, is steep. Recent methods for sampling IDPs by decomposing them into their component peptides and sampling these smaller systems individually before stitching them back together has been a fruitful approach in this space (Stelzl et al., 2022; Pietrek et al., 2020; Lindsay et al., 2021). However, this approach is limited, for example, in that it does not explicitly sample long distance interactions. The development of specific forcefields to accurately sample disordered proteins (Best et al., 2014; Song et al., 2017; Robustelli et al., 2018; Wu et al., 2018; Abascal and Vega, 2005) and enhanced sampling methods, like replica exchange with solute tempering (REST2) (Wang et al., 2011a), have also improved the tractibility of atomistic MD simulations of IDPs. However, efficiently computing properties of IDPs, especially on the condensate level, necessitates the use of coarse-grained models, either at the Martini level (Thomasen et al., 2022) or even more coarse grained at the single-bead-per-residue level (Regy et al., 2021; Tesei et al., 2021). Further levels of coarse graining have pushed the possible timescales and system sizes even further with lattice-models (Choi et al., 2019) and even machine-learning algorithms like ALBATROSS and CALVADOS (Lotthammer et al., 2024; Tesei and Lindorff-Larsen, 2022; von Bulow et al., 2024).

In this work, we employ computational techniques to understand how temperature impacts the structural character of the ELF3-PrD ensemble, how polyQ length and sequence context impacts these characteristics, and the impact of each of these factors on dynamics underlying ELF3 condensation. Using a fast and efficient chain-growth algorithm, we produce ensembles of ELF3-PrD at a variety of polyQ length and temperature conditions (Fig. 1B,1E). We discover temperature-responsive helices in polyQ-adjacent regions and identify a residue, F527, involved in the temperature-sensing mechanism of the PrD. Next, we employ all-atom replica exchange with solute tempering (REST2) simulations to characterize temperature-sensitive conformation changes that promote condensate formation (Fig. 1F). In addition to the wildtype ELF3-PrD, we examine the F527A mutant to characterize the key role played by the F527 residue in the temperature-response mechanism and a 0Q mutant with the variable polyQ tract removed to better understand the role of this polyQ tract in modulating the structural ensemble with increasing temperature. Finally, we perform coarse-grained Martini simulations to more directly study how ELF3-PrD condensation dynamics depend on temperature and polyQ length. Simulating 100 ELF3-PrD monomers in a small box allows us to investigate specific residues that play key roles in driving condensation, including a major role observed for aromatic residues in driving protein-protein interactions underlying condensate formation, while also revealing the impact of sequence context and the expansion or deletion of the variable polyQ tract on the temperature range, material properties and sensitivity of the temperature-responsive phase transition (Fig. 1G). Through this computational lens we are able to better understand the role and molecular mechanism of polyQ-adjacent helices and aromatic residues in the temperature-sensitive growth response of *Arabidopsis thaliana* mediated by the ELF3-PrD.

## Results

### Sequence-based insights on ELF3-PrD properties

Sequence heuristics were already initially useful to identify the existence of the ELF3-PrD (Jung et al., 2020; Lancaster et al., 2014), and further investigation of other heuristics can provide insight into its structural and dynamical properties. One simple first step is to look at the ratio of positive to negative charges in any IDP, which in the case of ELF3-PrD suggests a globular or ’tadpole-like’ shape (Fig. 1D) (Das and Pappu, 2013). Another simple sequence-level heuristic is the Ω_aro_, which indicates ‘patchiness’ of the aromatic residue distribution. For ELF3-PrD this value is high at 0.897, indicating the aromatics are distributed in clusters rather than evenly through the sequence (see Fig. S7 for a visualization of the aromatics within the ELF3-PrD sequence), which is in contrast to other well-known PrD’s like FUS that have a bias toward uniformly distributed aromatics (Martin et al., 2020). A further, sequence-based view is that of (Quiroz and Chilkoti, 2015), who demonstrated that repeats of Pro-*𝑋_𝑛_*-Gly were sufficient to provide a generic disordered scaffold that can be tuned to exhibit UCST or LCST characteristic phase separation in designed polymers. Inclusion of polar residues and zwitterionic pairs of charged residues in the X position or adjacent to the Pro-*𝑋_𝑛_*-Gly motifs, where n is ≤ 4, resulted in UCST character while LCST behavior results from the inclusion of nonpolar residues or lysine at these positions (see Fig. S7 for these motifs highlighted within the ELF3-PrD sequence). The *𝑋_𝑛_* component of these motifs is split between non-polar and polar residues with those proximal to aromatic residues being mostly non-polar, suggesting these regions may promote LCST phase separation.

Two sequence-based web utilities to quickly probe the structural features of IDPs are ALBA-TROSS and CALVADOS. ALBATROSS (Lotthammer et al., 2024) is a machine learning-based approach trained on coarse-grained simulations of synthetic disordered IDPs using Mpipi (Joseph et al., 2021b), a coarse-grained force field which emphasizes *𝜋*-*𝜋* interactions like those dominant in the dynamics of low-complexity domains (Vernon et al., 2018). CALVADOS (Tesei and Lindorff-Larsen, 2022) predicts structural properties based on an input sequence. However, CALVADOS does not utilize a deep learning algorithm, but instead runs a coarse-grained simulation in real time using a single bead-per-residue approach in which hydropathy scales inform amino acid interaction strengths. The Flory scaling exponent is calculated by ALBATROSS to be between 0.36 and 0.394 for each variant, slightly above the theoretical value for a collapsed globule, 0.33. This is in agreement with the globular or ’tadpole-like’ shape suggested by an earlier heuristic. The radius of gyration (*𝑅_𝑔_*) reported by ALBATROSS is 3.259 nm for 0Q, 7Q, and 19Q, with only the F527A mutant exhibiting a different value of 3.31, suggesting an insensitivity of ALBATROSS to variations in sequence. More structural predictions obtained from ALBATROSS for ELF3-PrD can be found in Table S1. Unlike ALBATROSS, CALVADOS was able to give estimates for values of interest for IDPs at different temperatures, however it estimated the WT Flory scaling exponent, *𝜈*, as 0.49, more suggestive of an ideal polymer with little self-interaction and a more extended shape, which is not in line with other methods. CALVADOS predicts a slightly larger *𝑅_𝑔_* value for the WT than ALBA-TROSS, with the 290K *𝑅_𝑔_* predicted as 3.41 nm. The WT ensemble is predicted to become more expanded at 340K with an *𝑅_𝑔_* of 3.73, as is the 13Q variant, shifting from 3.38 nm to 3.73 nm, the 19Q variant, moving from 3.7 nm to 3.8 nm, and the F527A mutant, expanding from an *𝑅_𝑔_* of 3.31 nm to 3.89 nm. Only the 0Q system is not predicted to undergo significant expansion upon heating from 290K to 340K. A comparison of ELF3-PrD *𝑅_𝑔_* values estimated with six different methods can be found in Fig. S8. A complete accounting of CALVADOS predictions for ELF3-PrD can be found in Table S2. While these quick approaches provide initial insights from just the amino acid sequence of any IDP, we found they sacrificed accuracy and precision in their ability to probe the range of environmental conditions relevant to characterize the effect of temperature and polyQ tract length on the conformational landscape of ELF3-PrD.

### Chain growth ensembles exhibit helicity in PolyQ-flanking residues

In order to understand how polyQ length and temperature change the behavior of ELF3-PrD, we examined the structural differences between ensembles with varying polyQ tract lengths constructed at different representative temperatures. Using the hierarchical chain growth (HCG) method by (Pietrek et al., 2020), we constructed ensembles of polyQ of 0, 7, 13 and 19 glutamine residues in length each at temperatures of 290K, 300K, 320K and 415K to produce a total of 16 ensembles with 32,000 conformations each, a total of 512,000 structures. These polyQ lengths were chosen to span deletion/short (0Q), WT-like (7Q), and expanded tracts (13Q/19Q), bracketing experimentally studied constructs and naturally occurring ELF3 polyQ polymorphism in Arabidopsis (Hutin et al., 2023; Jung et al., 2020; Undurraga Soledad et al., 2012). For each residue we summed the conformers with *𝛼*, 3_10_ or *𝜋*-helix character to obtain a total helical propensity (Fig. 1B). We found ELF3-PrD to be composed of short transient helices, with most having helical propensity of only around 5%. Though most of the ELF3-PrD exhibits only light helical character, the helices adjacent to the variable-length polyQ tract had significantly higher propensities, up to 30% for the N-terminal residues and 10% for the C-terminal residues. These regions, which we refer to as putative SLiMs, display some helicity with or without polyQ as seen in the 0Q system, but the polyQ tract seems to increase the portion of the ensemble with helical character (Totzeck et al., 2017). An additional feature of the polyQ-adjacent SLiMs is a degree of temperature-sensitivity not seen elsewhere in the PrD. In the 7Q system, the N-terminal polyQ adjacent SLiM falls from 30% at 290K to 20% at 300K and finally 10% propensity at 415K, while the same region in the 0Q system shows no appreciable reduction in propensity until 415K. The increase in temperature-responsiveness of these SLiMs in 7Q could be a by-product of the higher possible helical propensity enabled by the polyQ tract. Ensembles with polyQ tracts of 13 and 19 residues showed nearly identical results to the 7Q system (Fig. S9). This could be due to a limitation of the 5-residue fragments simulated to build libraries for the HCG method. Overall, it is clear the presence of a polyQ tract has a significant influence on the protein’s local structural character and temperature responsiveness.

E-PCA (Lindsay et al., 2018) is a contact analysis method that isolates the top sources of conformational variance (or dynamics when used on MD trajectories) by identifying residue pairs that form and break in a concerted manner. We apply it here to our HCG-generated ELF3-PrD ensembles to identify regions of correlated contacts which may be of mechanistic importance. For each system, the biggest signal consistently comes from three SLiMs directly N-terminally adjacent to the polyQ tract, suggesting correlated contact formation or breaking of contacts in this region explains most of the conformational variance, at least partially due to helix formation. At 290K, PC1 of all polyQ-containing systems (Fig. S10A, top-left panel, blue) contains a bimodal signal where each peak corresponds to interactions of residues within a polyQ-adjacent SLiM, suggesting some level of communication between these regions. The dominant mode of the 0Q system, on the other hand, exhibits a single peak, indicating communication between these SLiMs plays a less significant role in the contact dynamics of this system. At the maximum temperature, 415K, the contact dynamics of all systems converge, regardless of polyQ presence or length, as evidenced by the nearly identical PC1 values at this temperature (Fig. S10A, top-right panel, red). This peak at the high-temperature condition indicates formation of contacts that comprise the third polyQ N-terminal SLiM. However, one prominent negative value is consistently observed indicating interaction between F527 and Y530 strongly opposes contact formation within this SLiM. The persistence of this signal across ensembles and its location in the region identified as the primary source of conformational variability suggests a potential role for this contact pair in the general mechanism of ELF3-PrD, warranting further study. Further analysis of these chain growth ensembles can be found in the Supplementary Information.

### All-atom REST2 simulations reveal temperature-responsive conformation change in ELF3-PrD

While the static conformers generated by the HCG approach provided a relatively quick and resource-inexpensive look at the effects of varying the polyQ tract length and temperature on the local structure of the ELF3-PrD, we turned to a more intensive MD simulation approach to gain a more complete understanding of dynamics underlying ELF3 function. Based on trends seen in the initial HCG results, we decided to run three separate all-atom MD simulations using the REST2 method, including the wild-type ELF3-PrD to obtain a better understanding of the general molecular mechanism underlying temperature-responsiveness, the F527A mutant to understand the contribution of F527 to condensate formation and temperature responsiveness, and the 0Q system to isolate the role of the variable polyQ tract in ELF3-PrD dynamics. As with the HCG ensembles, helical propensity was calculated for all three PrD variants at four separate effective temperatures, 290K, 300K, 320K, and 405K (Fig. 2A, B and C respectively). Measurements of helical propensity around the variable-length polyQ tract from these trajectories are similar in trend to the HCG ensemble but with significantly larger propensity values, in the range of 30-65% instead of the 10-30%, for the N-terminal adjacent helix (Fig. 2A). Additionally, some level of helical enhancement was observed adjacent to all three polyQ tracts of ELF3-PrD rather than only the first as in the HCG case (Totzeck et al., 2017). The second polyQ tract (WT residues 568 to 572) exhibits enhancement in propensity and temperature-responsiveness in its N-terminal SLiM, and the third and last polyQ tract (WT residues 581 to 586) sees minor enhancement in its C-terminal SLiM, increasing to 10% or about double the baseline propensity. This enhancement of helical propensity indicates long-range interactions absent in the HCG ensembles further promote transient structure in the polyQ-flanking residues. The concerted nature of the enhancement of helicity and temperature-responsiveness may be more than a coincidence as enhancement of potential helical propensity could increase the ability to sense temperatures by providing a greater range of signals with more differentiable outcomes. The helical enhancement of polyQ-adjacent regions demonstrates how structural outcomes depend on sequence context, as degree and location of enhancement vary among the three tracts. The impact of sequence is not just local, as is apparent from the structural analysis of the F527A system. Relative to the WT, the F527A mutation attenuates helices, including those around each polyQ tract, despite being 15+ residues from the nearest tract.

**Figure 2.**
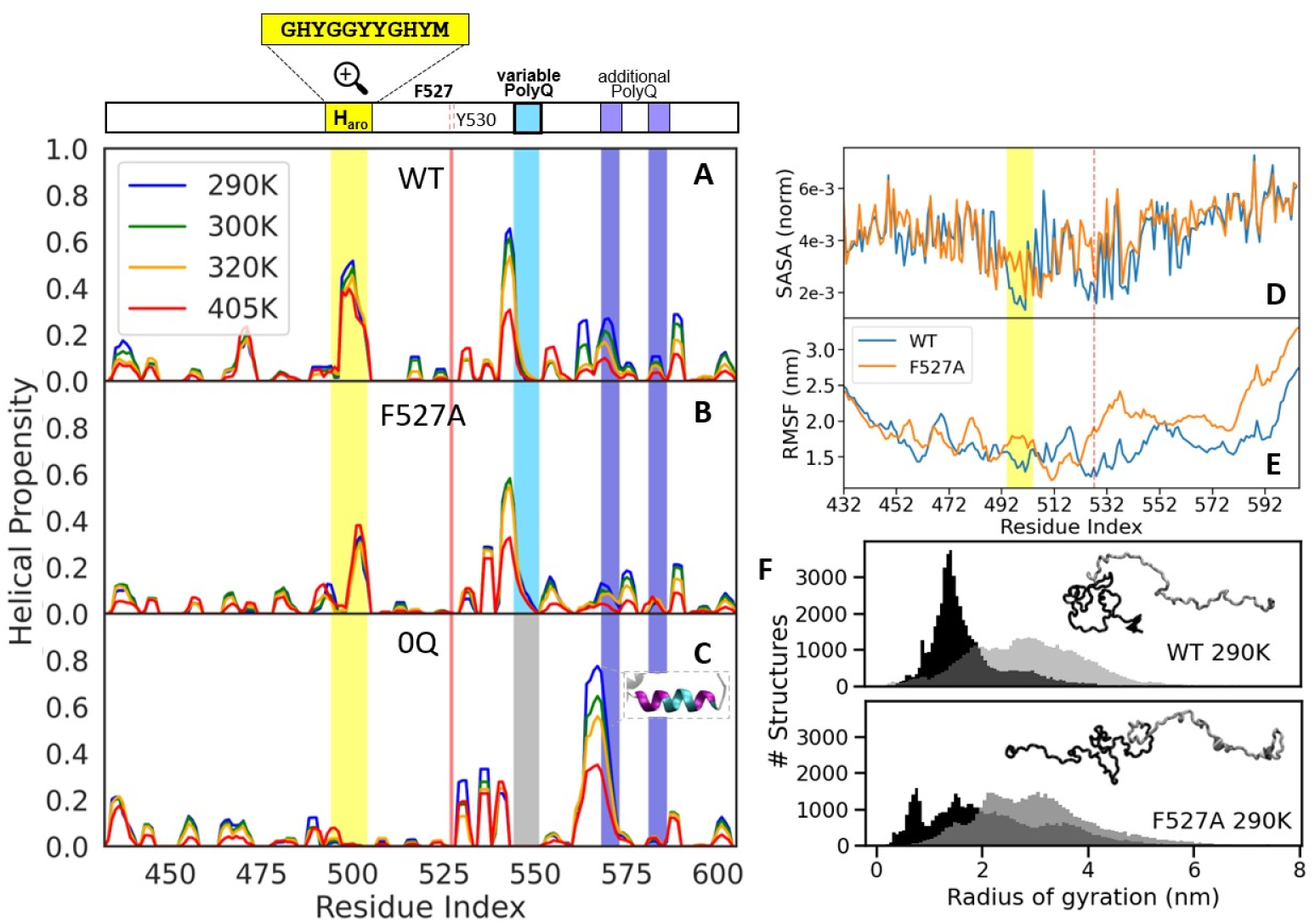
REST2 simulations reveal local and global conformation changes in response to temperature. A. Helical propensity of WT REST2 simulation at a range of temperatures. The variable polyQ tract is shaded in blue and the two polyQ tracts of consistent length are shaded in purple. A lengend denoting regions of interest is located at the top. B. Helical propensity of F527A REST2 simulation at a range of temperatures. The red line indicates the position of residue 527. C. Helical propensity of 0Q REST2 simulation at four temperatures. The gray shaded region indicates the removal of the variable polyQ tract. D. Top: Solvent-accessible surface area of WT and F527A systems taken from REST2 trajectories. E. The radius of gyration of the first 100 residues (black) and last 73 residues (grey) of ELF3-PrD. Top panel depicts and the WT and the bottom F527A, both at 290K.

While the 0Q system is reported to be significantly less sensitive to changes in temperature, some temperature-responsiveness does remain intact in experiments. The analysis of helical propensity for this trajectory reveals drastic structural deviation due to the absence of the polyQ (Fig. 5A), pointing to fundamental differences in the temperature-sensing mechanism compared with WT. In the absence of the variable polyQ tract, a large helix appears between residues 554 and 566 (Fig. 2C). This helix exhibits impressive stability at 290K (77% helicity) and rapidly declines linearly as temperature increases, bottoming out at 35% at 405K. This 0Q-native helix includes the first (second in WT) polyQ tract with the bulk of the helix forming N-terminal to polyQ. Aside from formation of this new helix, a distal aromatic helix, discussed further in the next section, is abolished in the 0Q system and helices that were flanking each side of the removed variable polyQ are attenuated. This result indicates the 0Q system may respond to temperature change by a mechanism vastly different from that of the WT, and the location of this new helix is further evidence that polyQ tracts promote helix formation in adjacent residues. Regardless, the interplay of variables like tract length (Totzeck et al., 2017), sequence context (Ramazzotti et al., 2012), tract location, and proximity to other polyQ tracts could all affect protein character and warrant further study.

### Aromatics play a major role in temperature-sensitive conformational change

The REST2 simulations painted a more complete picture of the ELF3-PrD dynamics and revealed some mechanistically important features absent in the HCG ensembles, partly due to the inclusion of long-range interactions. The most prominent was a 10-residue helix of heavy aromatic character (residues 494 to 504, “*𝐻_𝑎𝑟𝑜_*”) which was formed between 40-52% of the time in the REST2 WT simulations (Fig. 2A) but only present at baseline levels in the HCG ensembles. The F527A mutant sees a significant reduction in the size of this helix (Fig. 2B), suggesting long-range interactions with F527 may stabilize *𝐻_𝑎𝑟𝑜_*. Indeed, it seems this interaction may have consequences for the global conformation of ELF3-PrD. In the WT, both the regions around F527 and *𝐻_𝑎𝑟𝑜_* are largely buried, as shown by the low solvent-accessible surface area (SASA) in Fig. 2D, and their relative immobility as exhibited by RMSF values shown in Fig. 2E. A methionine residue of *𝐻_𝑎𝑟𝑜_*, M504, promotes compaction of this region through interaction of the sulfur atom with the ring of multiple local aromatic (tyrosine) residues (Fig. S11), a common but often overlooked motif (Valley et al., 2012). Upon mutation of F527 to alanine, both the F527 and *𝐻_𝑎𝑟𝑜_* regions become relatively solvent-exposed and flexible. This interaction influences the global shape of the protein by pushing the N-terminal region towards a globular conformation while the C-terminal region remains relatively extended. This is demonstrated by comparing the *𝑅_𝑔_* of the first 100 and last 73 residues for the WT and mutant systems, as in Fig. 2F. The first 100 residues of the WT (top panel) have a sharp peak indicating a compactness not observed in the mutant (bottom panel) while the last 73 residues sample a wide range of *𝑅_𝑔_* values in each case, indicating a more expanded and flexible nature. The mean *𝑅_𝑔_* of the whole PrD for the WT 290K ensemble is 2.75 nm, significantly more compact than predicted by heuristic models ALBATROSS and CALVADOS which reported values of 3.26 nm and 3.41 nm, respectively, as shown in Fig. S8.

Upon discovering the global structural influence of the long-range interaction between F527 and *𝐻_𝑎𝑟𝑜_*, we set out to determine its role in temperature-responsiveness. The minimum distances between atoms of F527 and *𝐻_𝑎𝑟𝑜_* were examined under different temperature conditions, and we found a linear increase in F527 and *𝐻_𝑎𝑟𝑜_* distance (and decrease in interaction) with increasing temperature, as demonstrated by the histograms in Fig. 3A. To get a better sense for the role of this temperature-responsive property in the conformational landscape of the protein, we plotted the F527-*𝐻_𝑎𝑟𝑜_* distance vs the radius of gyration in Fig. 3B. At 290K, the WT is largely homogeneous, mostly constrained to a single population of tightly interacting F527-*𝐻_𝑎𝑟𝑜_* and compact structures. However, with increasing temperature the ensemble shifts towards a second population that is slightly expanded with the F527-*𝐻_𝑎𝑟𝑜_* contacts broken. Representative structures of the low-temperature and emergent high-temperature populations are depicted in Fig. 3A on the left and right, respectively. The F527A mutant reinforces the importance of F527 in the WT conformational landscape as F527A samples a diverse range of populations in this space. However, the most persistent F527A population has significant overlap with the WT high temperature population, suggesting the F527A mutant may mimic the phenotype of WT ELF3-PrD at high temperatures. In Fig. 3C, we’ve employed the ELViM method (Viegas et al., 2024) to project the WT and F527A conformations onto a common space, where each conformer is a dot colored by the minimum distance between F527 and *𝐻_𝑎𝑟𝑜_* value. The projections on the left of Fig. 3C represent conformers from all four temperature conditions while those to the right represent specific temperature conditions. These projections further demonstrate the degree to which the F527A mutant alters the conformations sampled by ELF3-PrD and illustrates how increasing the temperature from 290K to 320K promotes access of the WT to new regions of conformational space. Additional visualizations and analysis of the ELF3-PrD conformation space can be found in the SI. Interestingly, in the 0Q system, the F527-*𝐻_𝑎𝑟𝑜_* interaction losses this temperature-responsiveness (Fig. S12) at all but the highest temperatures where we actually see an increase in the interaction propensity. This along with the 0Q-specific temperature-responsive helix suggest a different temperature-sensing mechanism in absence of the variable polyQ tract.

**Figure 3.**
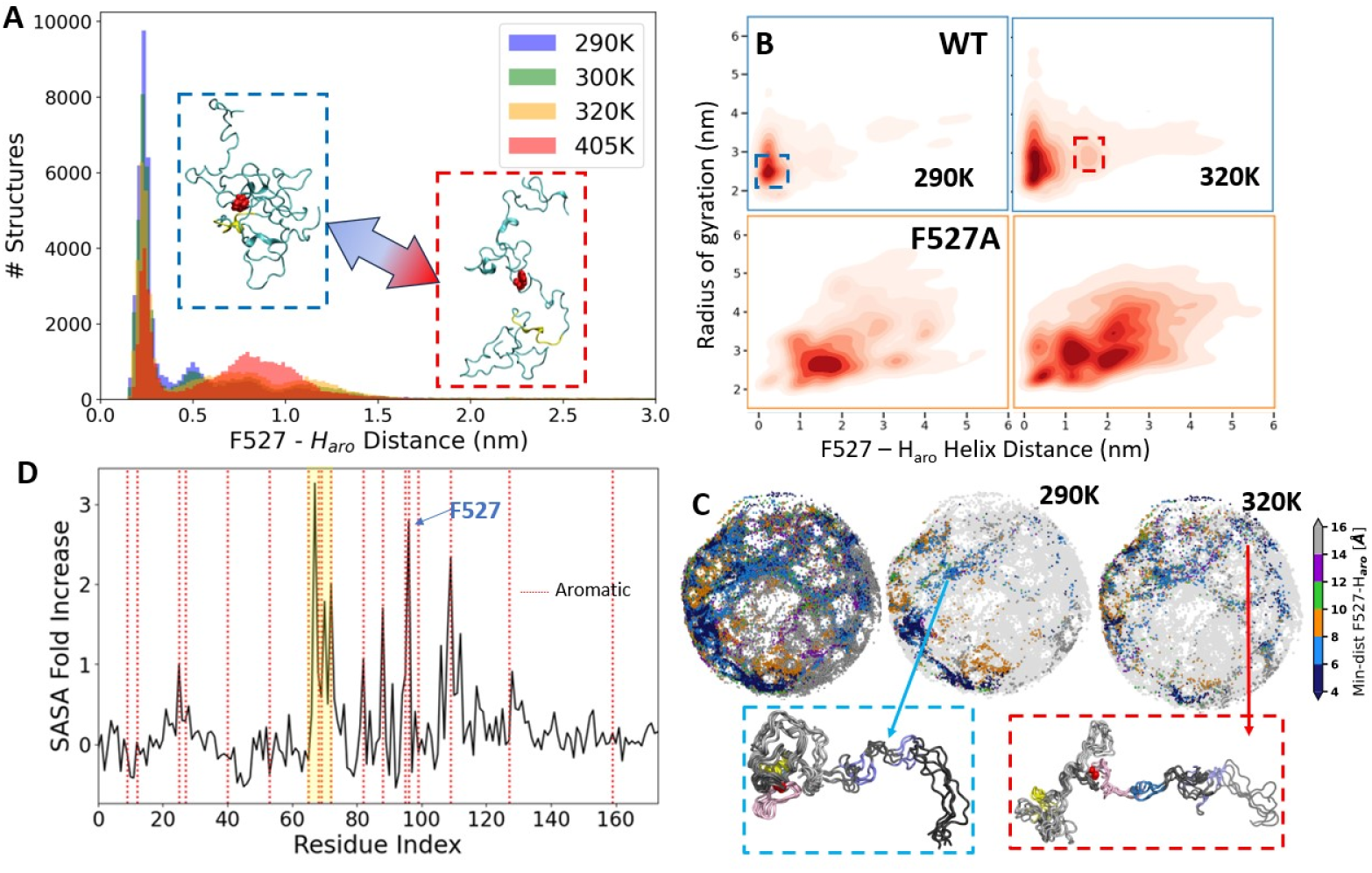
Temperature-responsive breaking of aromatic interactions exposes sticky aromatic residues to the solvent. A. The minimum distance between residue F527 and *𝐻_𝑎𝑟𝑜_* is plotted by temperature from the WT REST2 simulation. Representative low and high temperature structures are inside of a dashed blue or red box, respectively. F527 is in red and *𝐻_𝑎𝑟𝑜_* is in yellow. B. Free energy landscapes of WT (left) and F527A (right) with the X-axis defined as the minimum distance between residue 527 and *𝐻_𝑎𝑟𝑜_* and the y-axis defined as the radius of gyration. Each ensemble is represented at 290K and 320K. The blue box is the most populated region of conformation space at low temperature and the red box demarcates the emergent high temperature cluster. C. ELViM plots illustrating the conformation space of the full ensemble (left), the 290K ensemble (middle) and the 320K ensemble (right) where the color of each dot represents the minimum distance between F527 and *𝐻_𝑎𝑟𝑜_*. Below are example clusters of conformers with the left representing a commonly populated area at low temperature and to the right a cluster only observed at increased temperature. D. Fold increase in SASA between the identified lowand high-temperature populations. Dotted red lines represent aromatic residues and the yellow-shaded region are residues of *𝐻_𝑎𝑟𝑜_*.

The temperature-dependent breaking of the F527/*𝐻_𝑎𝑟𝑜_* interaction is particularly interesting due to its potential relevance to ELF3-PrD condensate formation. A significant increase in total SASA is observed in the high-temperature population with a major source being aromatic residues (Fig. S13). As previously mentioned, *𝐻_𝑎𝑟𝑜_* and F527 are regions of enhanced aromatic residue density, as seen from the dashed red lines in Fig. 3D. Breaking the interaction between F527/*𝐻_𝑎𝑟𝑜_* frees up 5 aromatic residues from these regions (Y496, Y499, Y500, Y503 & F527) and as well as four aromatic residues located between F527 and *𝐻_𝑎𝑟𝑜_* which can become buried during their interaction. Indeed, our high-temperature population exhibits enhanced solvent accessible surface area for these residues and for the majority of the 18 aromatic amino acids present in the ELF3-PrD with those residues located between *𝐻_𝑎𝑟𝑜_* and F527 especially affected (Fig. 3D). Given the well-established role of aromatic residues in driving interactions in low complexity disordered protein regions (Martin and Mittag, 2018a), it is reasonable to expect an increase in condensate formation propensity when these residues are accessible by other ELF3-PrD monomers. Indeed, previous work has established a link between single-chain dimensions and phase-separation propensity as a function of aromatic character (Dignon et al., 2018; Martin et al., 2020).

PolyQ solvation dynamics contribute to ELF3-PrD condensate forming propensity

A key question to understand temperature sensing by ELF3 is how expansion of the polyQ tract enhances the phase-separation propensity of ELF3-PrD, thus increasing the temperature sensitivity of condensation (Jung et al., 2020). One potential driver is the increasing availability of glutamine on the protein surface with increasing tract length. Plotting the SASA values for our HCG ensembles (Fig. 4A) highlights all three polyQ tracts as regions of persistent high solvent accessibility. While the SASA value of polyQ tracts is only moderately higher per residue than the average, indicated by a dashed black line, it persists for the length of the tract, creating patches of high solvent accessibility not seen elsewhere in the PrD. SASA values for these HCG ensembles were temperature-invariant for each case. Similar to the effect of increasing temperature in our REST2 simulations, increasing polyQ length enhances the overall solvent accessibility of the protein. PolyQ tract expansion provides a direct and tunable means to enhance PrD solvent-accessibility.

**Figure 4.**
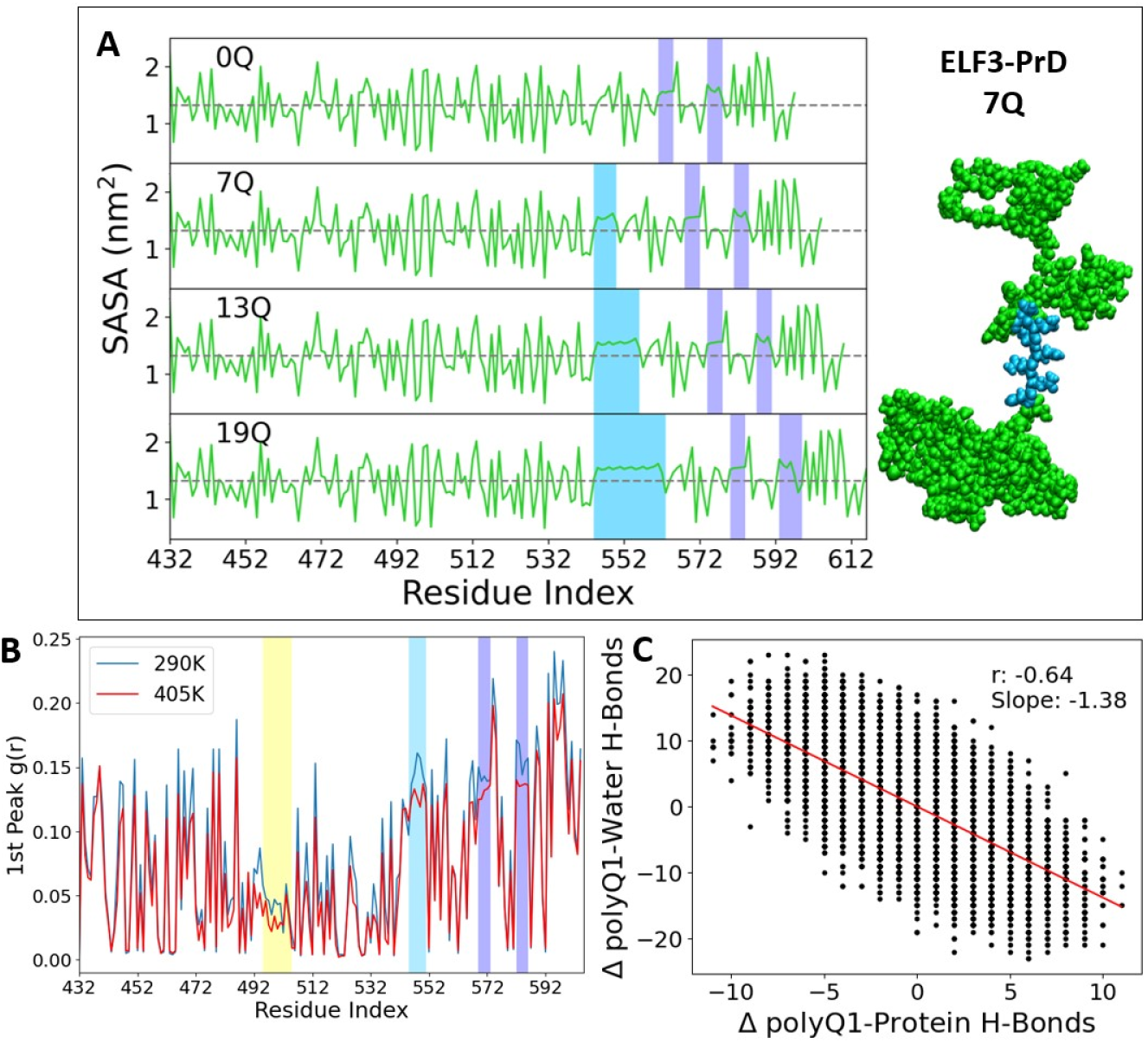
Cluster formation and monomer conformation change are driven by similar aromatic residues. A. The solvent-accessible surface area is shown for each HCG polyQ condition at 290K. The variable polyQ tract is highlighted in sky blue and the second and third polyQ tracts in a lighter blue. The black dashed line demarcates the average SASA value for the system. B. Per-residue hydration plot of ELF3-PrD WT at 290K (blue) and 405K (red). Each value represents the maximum height of the first peak of an RDF between the given residue and solvent oxygen atoms. The aromatic helix, *𝐻_𝑎𝑟𝑜_*, is highlighted in yellow and the polyQ tracts in sky blue. C. A scatter plot relating change in hydrogen bonds between the variable polyQ tract and water with the change in hydrogen bonds between polyQ and protein from the wild-type REST2 simulation at 290K. Breaking of polyQ-water hydrogen bonds is correlated with an increase in polyQ-protein bonds.

**Figure 5.**
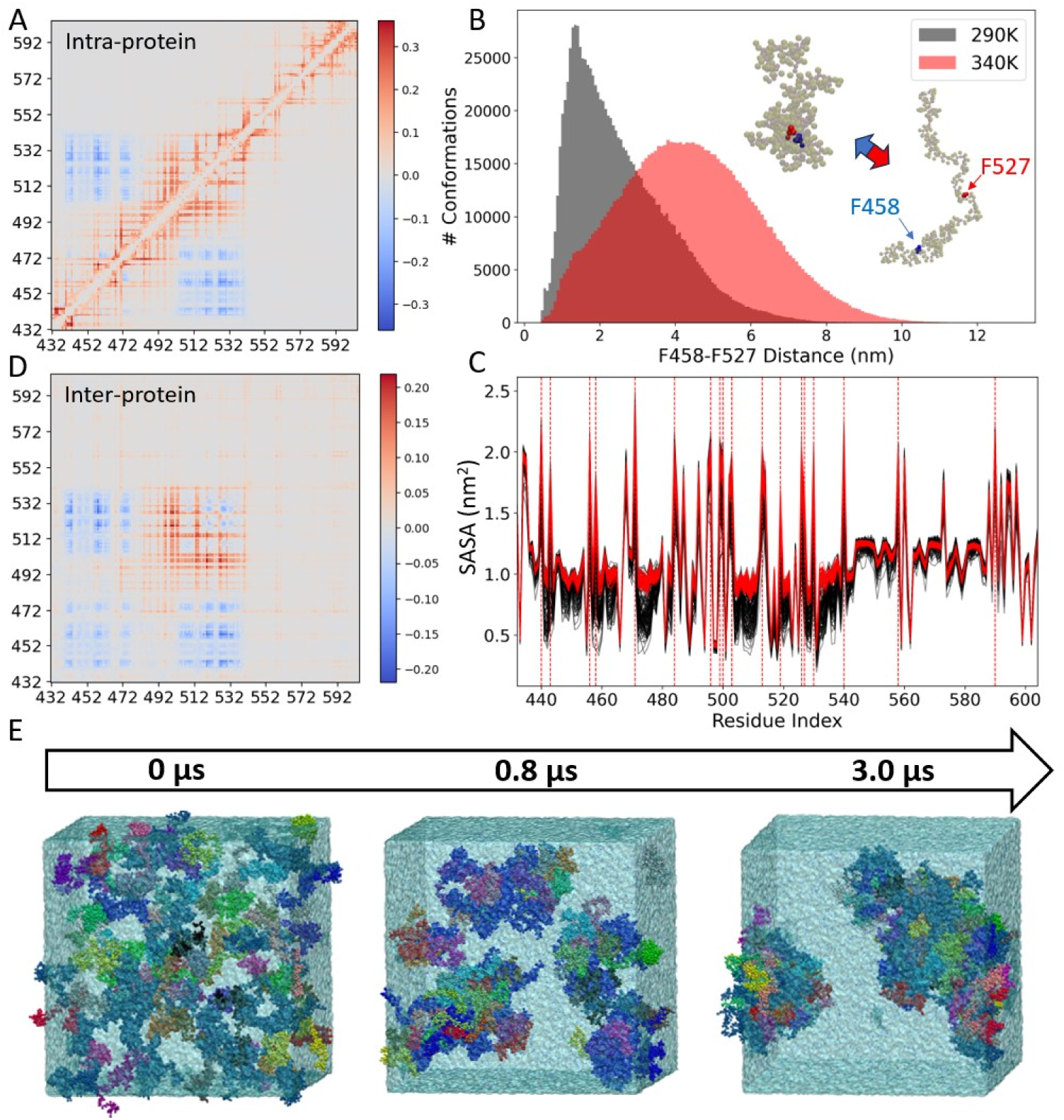
Temperature-dependent conformation change enhances SASA of CG monomers and promotes cluster formation driven by central aromatic residues. **A.** The intra-protein contact difference map between the 340K and 290K WT Martini condensate simulations. Values in red represent increases in contact value at 340K relative to 290K and blue represents a decrease. The entire 3 *𝜇*s trajectories were used to create these difference maps. **B.** Distance distributions between residue F527 and F458 at 290K (grey) and 340K (red) from WT CG condensate simulations. Distances are taken from intra-protein distances of each monomer. **C.** The solvent accessible surface area is plotted for all 100 monomers of both the 290K WT condensate simulation (black) and the 340K WT condensate simulation (red). **D.** The inter-protein contact difference map between the 340K and 290K WT Martini condensate simulations. As with Fig. 5A, contacts enhanced at high temperature are in red with those broken in blue. **E.** An illustration of the typical time evolution of an ELF3-PrD condensate simulation at 300K with simulation snapshots representing approximately 0 ns, 800 ns, and 3*𝜇*s, respectively.

Solvation dynamics serve as a determining factor in condensate formation behavior, in part by determining which regions remain accessible to the solvent and other proteins, and changes in temperature can alter these dynamics. To characterize how hydration differs across ELF3-PrD we obtained radial distribution functions (RDFs) of water oxygen atoms around ELF3-PrD as a whole and around each individual residue. Values for a per-residue solvation plot were obtained by using each residue as the reference point for an RDF calculation of water oxygen atoms and plotting the height of the first peak, representing the most tightly bound water molecules, referred to as the first solvation shell (Fig. 4B). At 405K, many residues exhibit a decrease in local solvation, with the most prominent and continuous decreases seen in the three polyQ tracts. Further investigation of ELF3-PrD polyQ interactions at high temperature show a tradeoff wherein a decrease in polyQwater hydrogen bonding is compensated by an increase in protein-protein hydrogen bonds, even though the number of polyQ-proximal water molecules increases (Fig. 4C). This is consistent with studies that have suggested water is a poor solvent for glutamine (Khare et al., 2005). This characteristic may influence condensation, either by promoting polyQ-protein interactions or altering the solubility of the protein.

### PolyQ tract length modulates temperature-responsiveness of PrD clustering

We next shift our focus from the structural ensembles of ELF3-PrD monomers to the condensation dynamics of the system. Performing coarse-grained Martini simulations provides a more direct way to identify what drives protein-protein interactions of the PrD and to observe the ways in which variables like polyQ tract length and sequence mutations modulate the formation and stability of clusters in response to temperature. We first examine the impact of temperature on contact dynamics of the 100 monomers that comprise our condensate simulations to see how they align with our all-atom simulations. The contact difference map seen in Fig. 5A provides an understanding of how intra-protein interactions change with increasing temperature. The blue regions indicate interactions between an N-terminal region of the PrD with centrally located aromatics at 290K, which are broken at 340K. Corresponding red lines in the central region indicate an enhancement of local interactions between aromatic residues within this region. This breaking of long-range contacts at increased temperature is similar to the temperature-induced conformational shift observed in the all-atom simulations. Here, F527 interacts in a temperature-dependent manner with F458 instead of the *𝐻_𝑎𝑟𝑜_* region (Fig. 5B), as was observed in the REST2 simulations. These Martini simulations do not capture transient helices, which may account for the lack of interaction with *𝐻_𝑎𝑟𝑜_*, but the trend is very similar nonetheless. By plotting the per-residue SASA of each of the 100 PrD monomers in Fig. 5C, a clear increase is seen across all residues, with the biggest peaks aligning with the aromatic residues. Based on a similar observation in the all-atom systems, we inferred that these residues would participate in inter-protein interactions, however, here we can observe these interactions directly. The inter-protein contact difference map in Fig. 5D indicate an increase in interaction between centrally-located aromatics at 340K as compared to 290K. An illustration of a WT PrD cluster forming at increased temperature is seen in Fig. 5E. The intra- and inter-protein difference maps are strikingly similar, suggesting the same interactions are responsible for monomer dynamics drive protein-protein interactions. We further explore the intra-protein interaction propensities of the aromatic residues in Fig. S14 that demonstrate the degree to which interactions between aromatic residues control the dynamics of this low complexity domain. The overall shape of the interaction profiles are nearly identical with only their scales varying, suggestive of the universal nature of aromatic contacts in these types of proteins. This plot provides some insight into the similarities observed in the intra- and inter-protein contact maps.

Our coarse-grained Martini simulations enable a direct characterization of how ELF3-PrD condensation depends on temperature, polyQ tract length, and sequence context. Following established approaches in protein condensate simulations, we quantify clustering using the size of the largest cluster as a proxy for condensation propensity (Benayad et al., 2021; Joseph et al., 2021a), with cluster membership defined by a 0.5 nm cutoff between backbone beads of distinct monomers. Starting from 100 monomers, each system rapidly coarsens into fewer, larger assemblies: the number of clusters decreases steeply over the first microsecond and then approaches a late-time plateau, consistent with progression toward a small number of dominant condensed assemblies (Fig. 6A). To assess temperature dependence, we compare the largest-cluster size across conditions for each construct (Fig. 6B, S17). For the wild-type (7Q) system, the dominant cluster increases markedly from 290K to 300K, a biologically relevant interval close to temperatures used in prior experiments on ELF3 thermoresponsiveness (Jung et al., 2020). Robust condensation persists through 320K, whereas at 340K the dominant cluster is strongly destabilized and approaches the condensation threshold (dashed line in Fig. 6B), indicating reduced persistence of the condensed phase at high temperature.

**Figure 6.**
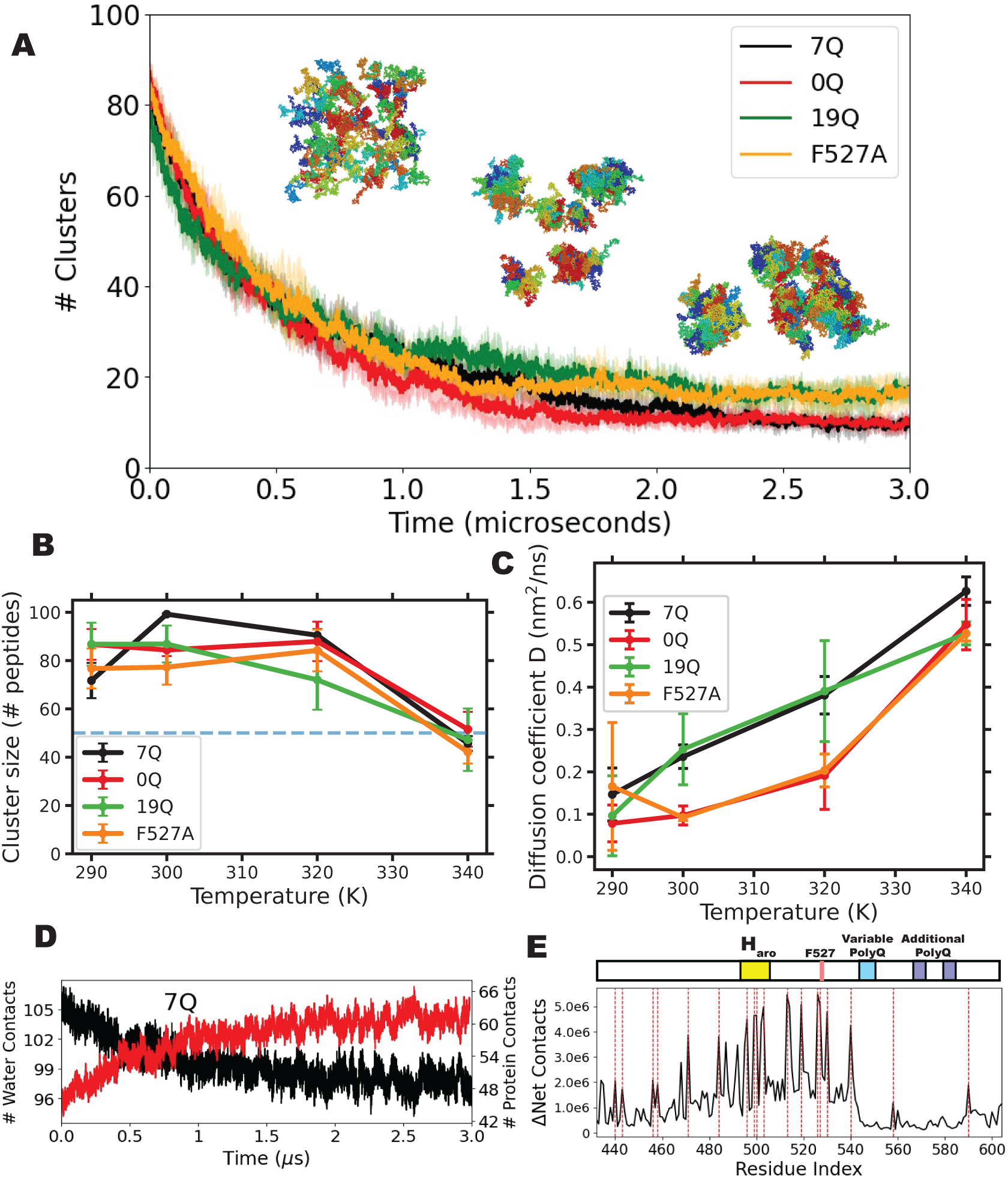
Martini 3 simulations reveal temperature- and variant-dependent condensate stability and internal dynamics of ELF3-PrD. Please check “Analysis of Martini Condensate Simulations” for more details of the methods A. Time evolution of the # of clusters at 290 K for 7Q (black), 0Q (red), 19Q (green), and F527A (orange); shaded regions show variability across independent replicas. Representative simulation snapshots illustrate coarsening from many small clusters to fewer larger assemblies. B. Condensate stability quantified as the mean size of the largest cluster (number of peptides) as a function of temperature for each variant (mean ± SEM calculated over the last 1 *𝜇𝑠* of three independent replicas). The dashed line indicates the threshold used to define a condensed state (*𝑁*_max_ *>* 50). C. Internal mobility within the condensed phase quantified by the effective diffusion coefficient *𝐷*extracted from peptide mean-squared displacement analysis (mean ± SEM across three independent replicas), shown as a function of temperature for each variant. D. Desolvation of the variable polyQ region during condensation illustrated for 7Q at 300 K: the number of water contacts to the polyQ tract decreases (black) as the number of protein contacts increases (red) over time. E. Residue-resolved change in inter-protein contacts for wild-type (7Q) at 300 K relative to 290 K (ΔNet Contacts), highlighting regions spanning the aromatic helix (*𝐻*_aro_), F527, and the variable/additional polyQ segments. Aromatic residues are marked by dashed red lines.

PolyQ length and sequence context tune not only condensate stability but also internal dynamics within the condensed phase. Across all constructs, effective internal diffusivity increases with temperature (Fig. 6C, S19, S20), indicating faster internal rearrangements at higher temperature. However, peptides can remain similarly condensed while exhibiting distinct mobility: for example, 0Q forms large dominant clusters over 290–320K (Fig. 6B, S17) yet exhibits reduced internal mobility relative to 19Q (Fig. 6C), implying that polyQ variation can modulate the material state of the condensate. Complementary analyses of anomalous diffusion (*𝛼*) and self van Hove displacement distributions (Fig. S19-21) further support this interpretation by indicating persistent subdiffusion and dynamic heterogeneity within the condensed phase. Consistent with this, shape anisotropy (*𝜅*^2^) and radius-of-gyration (*𝑅_𝑔_*) distributions at 300K largely overlap across variants, with *𝜅*^2^ spanning nearly the full 0–1 range, indicating broad morphological heterogeneity (Fig. S22–S23). To connect these mesoscale behaviors to molecular drivers, we examined desolvation and residue-resolved contact formation during condensation. As clusters mature, protein–water contacts decrease while protein–protein contacts increase, consistent with progressive desolvation coupled to strengthening intermolecular interactions (Fig. 6D). A residue-resolved analysis of temperature-dependent contact changes shows that the net increase in contacts at 300K compared to 290K is not uniform along the sequence: the largest gains localize to aromatic-rich regions, including a prominent hotspot near the F527-adjacent segment (Fig. 6E), consistent with a sticker-dominated contact network that strengthens with temperature.

PolyQ expansion and deletion reshape these temperature dependencies in distinct ways. Expansion to 19Q supports robust condensation at 290–300K but yields reduced dominant-cluster size at 320K relative to 7Q (Fig. 6B, S17), indicating that longer polyQ does not uniformly enhance clustering across all temperatures. Notably, the polyQ tract in 19Q is less hydrated when normalized by tract length: the distribution of “waters per glutamine residue” is shifted to lower values for 19Q compared with 7Q (Fig. S16), consistent with a relative reduction in per-residue solvation of the expanded tract. Conversely, deletion of the tract (0Q) does not abolish condensation but alters the balance between condensate stability and internal dynamics (Fig. 6B,C,S18-21), supporting a primary role for polyQ length in tuning condensate material properties rather than acting as an all-or-none switch for clustering.

Finally, we probed sequence context using the F527A mutant. F527A forms a robust condensed phase over 290–320K with modest shifts in largest-cluster size (Fig. 6B, S17), but exhibits destabilization at 340K and altered internal mobility (Fig. 6C), consistent with disruption of a key aromatic contact hotspot identified in Fig. 6E. Together, these results support a mechanism in which temperature-responsive condensation is mediated by a sticker-rich contact network dominated by aromatic motifs, while polyQ length tunes the temperature dependence of condensate stability and the dynamical (material) state of the condensed phase.

## Discussion

We have used varying levels of simulation accuracy to understand the molecular basis for the ELF3-PrD-based temperature response in plants. Together, these methods have enabled us to propose mechanisms by which ELF3-PrD responds to temperature to promote protein-protein interaction and condensation. Initial sequence-based methods allowed us to put the ELF3-PrD in the context of other IDPs: a ‘tadpole’-like weak poly-ampholite (Mao et al., 2010) with some sequence characteristics in line with those seen for other LCST proteins (Martin and Mittag, 2018b; Dignon et al., 2019). Furthermore CALVADOS indicated some temperature dependent changes in radius of gyration, but CALVADOS and ALBATROSS gave different results for values like the Flory scaling exponent. However, we sought a more molecular-level understanding of the temperature-dependent properties of this process by building HCG ensembles at different temperatures and with different polyQ lengths to find that the variable polyQ was flanked by temperature-sensitive helices. These initial HCG results also indicated the importance of aromatic residues – specifically the F527. Extensive molecular simulations with REST2 allowed us to observe the temperature-dependent conformation landscape of the ELF3-PrD, including crucial long-range aromatic interactions not captured by HCG, like that of F527 and *𝐻_𝑎𝑟𝑜_*. Note that with an initial preprint the HCG and REST2 data of the atomistic ELF3-PrD simulations were made public at an early stage, and several groups analyzed these terabytes of data to test their methods, and found similar corroborating analyses to those presented here (Viegas et al., 2024; González-Delgado et al., 2024).

To connect these atomistic insights to collective condensate behavior, we turned to coarsegrained Martini simulations, used here primarily to assess comparative trends across constructs and temperatures. Coarse-grained simulations bridge temperature-dependent single-chain behavior of ELF3-PrD to emergent condensate properties and provide a mechanistic framework for interpreting how polyQ variation and sequence context tune condensation. This mesoscale picture is consistent with Hutin *et al*., who observed temperature-responsive condensation across Q0/Q7/Q20 constructs and emphasized that polyQ length modulates condensate properties and maturation rather than acting as a strict on/off determinant of phase separation (Hutin et al., 2023). In this framework, increasing temperature promotes net desolvation and increases the engagement of a sequence-localized cohesive network, with the largest temperature-dependent gains in intermolecular contacts concentrated in aromatic-rich segments, including the region adjacent to F527, consistent with an aromatic “sticker” network as a primary determinant of condensate cohesion.

The resulting condensed phase is not a simple Brownian liquid in this regime, but exhibits heterogeneous, viscoelastic dynamics: internal mobility increases with temperature, yet remains subdiffusive and dynamically heterogeneous (persistent *𝛼 <* 1 and non-Gaussian displacement statistics), consistent with caging-and-hopping-like rearrangements within the dense phase. This molecular picture aligns with Hutin *et al*.’s observation that ELF3-PrD condensates can rapidly age toward a hydrogel-like, low-recovery state while retaining temperature-tunable behavior (Hutin et al., 2023). Within a sticker–spacer interpretation, the variable polyQ tract functions as a tunable spacer that reshapes packing, hydration, and rearrangement kinetics with temperature, thereby modulating the stability and dynamical state of the condensate across a biologically relevant window. Disrupting a key aromatic sticker (F527A) perturbs this balance, underscoring that a small number of sequence-encoded motifs can exert disproportionate control over temperature-dependent condensate properties.

Overall, this molecular approach has helped understand the interplay of the varied sequence contexts of the ELF3-PrD and how they effect transient structures and temperature dependence. Here we found that aromatic residues, especially F527 and the central aromatic cluster, are key to condensate formation for this IDP, consistent with other analyses of condensate-forming IDPs (Martin et al., 2020). Furthermore, the transient temperature-dependent helices flanking the polyQ’s points to the importance of the sequence context around the polyQ’s. It is possible it is exactly this context that has reinforced these domains and the plasticity of polyQ motifs for adaptive temperature sensing, even when excessive polyQ length leads to disease (Reiner et al., 2011). These transient helices are reminiscent of SLiMs found in other IDP systems (Davey et al., 2012).While SLiMs have been seen to play a role in condensate formation, for example in TDP43, here we see a decrease in helicity as temperature increases, so this SLiM formation anti-correlates with condensate formation. The breakdown of these transient helices at high temperatures exposes aromatic residues to enhance condensate formation at higher temperatures (Fig. S27). Furthermore, a possible role for these putative SLiMS around the polyQ motifs is to enhance protein-protein and protein-DNA interaction with the rest of the EC complex at lower temperatures, when they are more stable.

Understanding the role of these transient helices in condensate formation from molecular simulation is complicated by the pros and cons of the various simulation methods used. The HCG and REST2 simulation both consistently found these polyQ-adjacent temperature-dependent helices, actually more prominent in the more accurate REST2 simulations (though we acknowledge reports of REST2 sampling overly compact states (Zhang and Chen, 2022)). These helices, however, are not well-captured in the coarse-grained simulations necessary to simulate condensate formation, as the only option is to explicitly enforce them with an elastic network model. However, we believe that the overall properties of these coarse-grained simulations are encouraging, as the contact maps and radius of gyration were overall very similar to those seen in the REST2 simulations. Furthermore, that Martini 3 was able to show increase in condensate-forming propensity at higher temperatures for the WT ELF3-PrD and an increase in condensate-forming propensity for a longer polyQ tract is encouraging for the relevance of this force-field for studying LCST systems. For example, our attempts at using other single-residue-per-bead forcefields with implicit solvent like HPS (Dignon et al., 2019) lead to much more compact conformations than we observed in atomistic simulation (Fig. S24), and especially noting the importance of protein-water interactions for LCST systems, we chose to not go this route.

The importance of understanding the ELF3-PrD temperature sensing mechanism in plants has increased since its original discovery (Jung et al., 2020) as more similar systems have been described (Zhao et al., 2017; Bohn et al., 2024), and the threats of a warming planet on plant and crop health have become ever clearer. The results we present here help elucidate some of the molecular origins of this temperature sensitivity, however it is clear there is much work still to do. For example the simplified system we have studied here of the 173 residue ELF-PrD without the rest of the 695 residue protein and in the absence of the members of the EC (LUX, ELF4, and DNA) begs for further studies within a more realistic context, though ELF3-PrD condensate formation has been seen *in vitro*. Relatedly, both *in vitro* and *in vivo* experimental validation of the molecular mechanisms we postulate is an essential path forward. Overall, though, we believe that this study has taken the cutting edge of computational capabilities in the field of biomolecular condensate toward understanding a biologically important LCST.

## Methods

### Hierarchical Chain Growth Method and Simulation Details

The sequence for *Arabidopsis thaliana* ELF3 was obtained from UniProt. The PrD sequence (residues 432 to 604) was isolated and divided into 57 segments. Each segment was five amino acids in length including two residues that overlap with the previous segment and two residues that overlap with the subsequent segment. Each of the 57 segments was capped with an N-terminal acetyl group and a C-terminal N-methyl group to neutralize interactions of the charged ends of the protein. Segments were parametrized using the Amber03ws forcefield (Best et al., 2014) and placed in a periodic rectangular box with 1nm of padding on each side of the protein and solvated with TIP4p2005 water molecules (Abascal and Vega, 2005).

Each segment was minimized until the maximum force reduction dropped below 1000 kJ/mol/nm per minimization step. For each of the 57 segments, 24 replicates were equilibrated to various temperatures ranging from 290K to 405K for 10ns. For this step, temperature effects were implemented using the v-rescale thermostat and the Berendsen barostat (Berendsen et al., 1984). Finally, replica exchange MD was run for each segment for 100ns using GROMACS version 2021.1 (Abraham et al., 2015). For this production run, the v-rescale thermostat was used to maintain temperature and the Parrinello-Rahman barostat ((Parrinello and Rahman, 1980)) was used to maintain a pressure of 1 bar.

Once the fragment simulations were finished, we used the hierarchical chain growth (HCG) application, developed by Pietrek *et al*., to construct a 7Q ELF3-PrD ensemble from these 57 segments consisting of 32,000 unique conformations (Pietrek et al., 2020; Stelzl et al., 2022). Conformers were randomly chosen from each segment and joined in a pairwise fashion to create a set of fragment dimers. These were then joined into tetramers, octamers, and so on until the full 57-segment ELF3-PrD was reconstructed. This process was repeated until we had obtained ensembles with polyQ tracts of 0, 7, 13 and 19 residues in length each at temperatures of 290K, 300K, 320K and 415K for a total of 16 ensembles. The selected tract lengths (0/7/13/19Q) were intended to bracket deletion/WT-like/expanded variants that are experimentally and biologically motivated (Hutin et al., 2023; Jung et al., 2020; Undurraga Soledad et al., 2012). Due to the two-residue overlap, we were required to run 19 segments made up of residues C-terminal to the polyQ region in order build the 0Q, 13Q and 19Q systems.

### REST2 Simulation Details

The HCG method enabled us to quickly look at large ensembles of different polyQ conditions for potential structural differences and pointed us towards some regions and residues of interest. The usefulness of the HCG method, for our purposes, is limited by the absence of long-range interactions. In order to accurately study the dynamics of ELF3-PrD we turn to REST2 (Replica Exchange with Solute Tempering), an all-atom enhanced sampling technique capable of exploring longer effective time scales than traditional all-atom MD (Jo and Jiang, 2015). Much like temperature replica exchange (T-REMD), REST2 runs multiple replicas in parallel, each at a different effective temperature. Every few hundred MD steps, replicas attempt to swap with one another with the probability of acceptance tied to the energy difference between replicates (Wang et al., 2011b). Unlike T-REMD where temperatures values are set for each replica, REST2 modulates the strength of protein-protein and protein-water interactions to achieve an effective temperature, a feature which reduces the number of replicas required compared with T-REMD. While REST2 can still be resource intensive, it is one of few methods able to provide Boltzmann-distributed ensembles of IDPs at biologically relevant timescales.

REST2 simulations (Wang et al., 2011b) were performed of the full-length wild-type ELF3-PrD, the ELF3-PrD F527A mutant and the 0Q ELF3-PrD with the variable polyQ tract removed. For the initial structure of these simulations, we chose an especially helical conformer from our HCG wild-type ensemble created at 290K. For the F527A mutant, F527 was manually edited to alanine by removing side-chain atoms and changing the residue name in the structure file. Each system was inserted into a rectangular periodic box with 1 nm of padding around each side of the protein and treated with the Amber03ws forcefield (Best et al., 2014) and the TIP4p2005 water model (Abascal and Vega, 2005). The wild-type system contained 2606 protein atoms, 122623 TIP4P water molecules, 439 Na atoms and 342 Cl atoms for a total of 490489 atoms. The F527A mutant matched this setup but with 2596 protein atoms for a total of 493769 atoms. The 0Q system contained 2487 protein atoms, 210607 water molecules, 237 Na atoms and 240 Cl atoms for a total of 846088 atoms. A 50ns equilibration run was performed for each system using the v-rescale thermostat and Berendsen barostat (Berendsen et al., 1984). After equilibration, each protein atom was marked within the base topology file as a “hot atom” and replicas were created by scaling interactions involving these “hot atoms” by a lambda value corresponding to temperatures in the range of 290K to 405K. The wild-type and F527A systems each had 20 replicas while 0Q had 12 replicas in order to achieve an exchange probability between 35% and 45%. REST2 production runs of each system were performed for 500ns using the v-rescale thermostat and Parrinello-Rahman barostat (Parrinello and Rahman, 1980). Convergence was verified using a split analysis of the radius of gyration values for each system at multiple temperatures (Fig. S25).

### Martini Simulation Details

All condensate simulations contain 100 ELF3-PrD monomers in a 40 nm^3^ box. Monomers used for the initial configurations were obtained by performing an RMSD-based cluster analysis on previously run all-atom REST2 simulations of ELF3-PrD monomer. A representative structure was taken from 10 of the most populated clusters and each was converted to Martini format using Martini 3.0 and inserted into the box 10 times for a total of 100 monomers. Protein-water interactions were scaled up by 10%, a procedure shown to bring simulated proteins into better agreement with experimental measurements (Thomasen et al., 2022). Each initial configuration was equilibrated for 50ns to four different temperatures, 290K, 300K, 320K and 340K followed by production runs of more than 3 *𝜇*s. This process was performed for wildtype (7Q), 0Q, and 19Q ELF3-PrD, as well as the F527A mutant. All systems used explicit Martini water and contained 150mM NaCl. Three replicates were performed for each condition.

### Analysis of Martini Condensate Simulations

Cluster identification and condensate stability.

Clusters were identified using a distance-based graph criterion: two monomers were considered connected if any pair of backbone beads (BB) between the monomers approached within 0.5 nm, and clusters were defined as connected components of this graph (Benayad et al., 2021; Joseph et al., 2021a). For each trajectory we tracked (i) the number of clusters and (ii) the size of the largest cluster *𝑁*_max_(*𝑡*) as primary condensation observables. To quantify condensate stability in a statistically robust manner, we computed the mean largest-cluster size over a late-time analysis window (final 1 *𝜇*s of each trajectory) and reported mean ± SEM across independent replicas. We also computed a condensate lifetime defined as the fraction of frames in which *𝑁*_max_(*𝑡*) exceeded a fixed threshold (*𝑁*_th_ = 50 peptides), again reporting mean ± SEM across replicas.

Condensate dynamics: internal mobility, anomalous diffusion, and van Hove analysis. To characterize internal mobility within the condensed phase, we computed the mean-squared displacement (MSD) of monomers (center-of-mass trajectories in the condensed phase as a function of lag time *𝜏*,

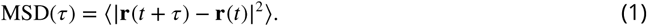

An effective diffusion coefficient *𝐷* was estimated from a short-time linear fit of MSD(*𝜏*) over *𝜏* ≤ 50 ns using MSD(*𝜏*) ≈ 6*𝐷𝜏*. To quantify deviations from normal diffusion, we computed an anomalous diffusion exponent *𝛼* by fitting MSD(*𝜏*) v *𝜏^𝛼^* on a log–log scale. Dynamic heterogeneity was assessed using the self part of the van Hove function evaluated at 1 ns lag time.

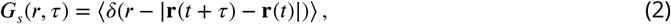

To highlight non-Gaussian behavior, *𝐺_𝑠_*(*𝑟, 𝜏*) was compared to a Gaussian reference distribution with variance matched to the MSD at the same *𝜏* (Supplementary Information).

Condensate morphology and chain compaction.

To quantify condensate morphology, we computed the shape anisotropy *𝜅*^2^ of the largest cluster from the eigenvalues *𝜆*_1_*, 𝜆*_2_*, 𝜆*_3_ of its gyration tensor,

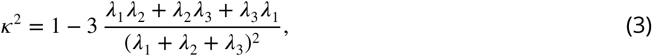

where *𝜅*^2^ = 0 corresponds to a spherical object and larger values indicate increasing asphericity. Distributions of *𝜅*^2^ were summarized using violin plots across frames and replicas. We also quantified intrachain compaction by computing per-monomer radii of gyration *𝑅_𝑔_* within the condensed phase and reporting the corresponding distributions (Supplementary Information). All coarse-grained analyses were performed using in-house Python scripts; replicate uncertainty was reported as mean ± SEM unless otherwise stated.

### Statistical Analysis of Ensemble Contact Maps

We used two types of contact analysis to identify mechanistically important interactions and characterize differences in our ELF3-PrD chain-growth ensembles. Both approaches, E-PCA and I-PCA (Johnson et al., 2018), are part of the CAMERRA family of contact analyses developed by Shen *et al*. and seek to explain sources of conformational variance of protein ensembles. In E-PCA, contact maps from each conformation of the ensemble are used to produce a contact covariance matrix of each possible residue pair, *𝑖*and *𝑗*, with all other residue pairs, *𝑘* and *𝑙*. PCA is performed using this covariance matrix as input and the top PCs are returned. This method has been used to uncover coordinated interactions underlying mechanisms of allostery, cooperative ligand binding, and general collective motions of a variety of proteins (Lindsay et al., 2018; Clark et al., 2016; Johnson et al., 2015a,b). Due to the absence of long-range interactions in the HCG method used to generate our ensembles, E-PCA signals will be more prominent for local interactions. The dynamics of folded proteins tend to be dominated by the top few modes (PCs), however, intrinsically disordered proteins exhibit a slower eigen decay in which the first several PCs may contribute nearly equally (Fig. S26).

The second CAMERRA variant, I-PCA, provides information on the packing of residues and can identify regions of the sequence in which residues tend to interact more frequently. In this approach, individual residue pairs are considered against the rest of the protein, resulting in a covariance matrix of size *𝑁*× *𝑁*as opposed to *𝑁*^2^ × *𝑁*^2^ for the E-PCA variant. This approach has been prominently used in studies of chromatin structure to differentiate transcriptionally active and inactive regions (euchromatin and heterochromatin, respectively) from Hi-C maps (Lajoie et al., 2015). When applied to natively structured proteins I-PCA tends to identify the individual domains of the protein and while IDPs do not contain structured domains, I-PCA can nevertheless identify dynamically associated domains (Lindsay et al., 2021).

## Acknowledgments

We would like to acknowledge Lucy Reading-Ikkanda for providing the illustrations in Fig. 1A, Lukas Stelzl and Lisa Pietrek for developing the hierarchical chain growth method used here, and Javier González Delgado and Juan Cortes for characterizing the conformational landscape of the ELF3 REST2 simulations using their WARIO contact analysis method and subsequent discussions. VBPL is supported by the Brazilian agencies FAPESP (Grants 2023/02219-1 and 2022/07231-7) and National Council for Scientific and Technological Development – CNPq (Grant 310017/2020-3).

## Data Availability

The datasets generated and analyzed during the current study are available for download: HCG ensembles: https://users.flatironinstitute.org/~ccb/smbp/elf3_data/elf3prd_hcg.zip REST2 trajectories: https://zenodo.org/doi/10.5281/zenodo.13226984 Martini trajectories: https://zenodo.org/doi/10.5281/zenodo.13285376 Analysis scripts and code: https://github.com/flatironinstitute/ELF3

