## Supplement for "Molecular dynamics simulations illuminate the role of sequence context in the ELF3-PrD-based temperature sensing mechanism in plants"

This document contains:

1. Supplementary Text
2. Supplementary Figures

### Supplementary Text

#### Supplementary Information

##### Potential Limitations of REST2, and MD in General, in the Study of IDPs

REST2 simulations were important for discerning the dynamics of ELF3-PrD in this study. While this method provides a way to sample more of the conformational landscape of IDPs than traditional all-atom MD or REMD simulations, there is some evidence that the REST2 protocol favors condensed states of IDPs over more extended states.[1] Additionally, MD force-fields are not typically tuned for optimal quantitative results at a wide range of temperatures. Nonetheless, qualitative trends can be ascertained. Some of the temperatures examined here are beyond what is likely to be observed in nature, however, with the limitations considered here in mind, we believe our results to be relevant at biological temperature ranges.

##### Contact Analysis of Chain Growth Ensembles with CAMERRA

While E-PCA can be informative about the concerted dynamics of individual pairs of residues, I-PCA can reveal the existence of modular domains and their spatial relationships, which can be especially useful for IDPs in which typical structured domains are not present. I-PCA was performed on each system to investigate how the polyQ tract and temperature affect which regions interact to drive the protein dynamics. The signal of PC1, like that of E-PCA, is focused almost exclusively on the region around the polyQ tract and the three N-terminally adjacent SLiMs (Fig. S1A), which are indicated here as the three prongs of the curve in PC1. The emphasis on these SLiMs is again an indicator they are largely responsible for the conformational variance of our HCG ensembles. In polyQ-containing systems, at 290K, PC1 indicates interactions between the residues of the polyQ N-terminal SLiM helix dominate the contact dynamics as the only region of dense residue packing. As the temperature rises, all three SLiMs N-terminal to polyQ increasingly interact to form a single spatially-associated domain, indicated by the leveling out of the three prongs in Fig. S1A. This high-temperature result of 7Q matches what is consistently observed in PC1 of the 0Q system regardless of temperature. These results suggest enhancement of the N-terminal polyQ-adjacent SLiM accounts for the differentiation between temperature conditions observed in PC1, whereas the 0Q system in which this enhancement is not observed demonstrates a consistent signal nearly invariant of temperature.

We next examine the second PC of the I-PCA analysis of ELF3-PrD, which accounts for nearly the same amount of variance as PC1, as evidenced as by the slow eigenvalue decay (Fig. S16). PC2 of 7Q (Fig. S1C) indicates two distinct domains are present at 290K. The domain illustrated in red in the illustration in Fig. S1C is comprised of residues of, and N-terminal to, the third SLiM from the polyQ tract as well as a region between residues ~491 and 501, all of which exhibit negative PC2 values. This region is comprised largely of aromatic residues and is referred to here as "$H_{aro}$". Residue F527 appears again here as the residue with the most negative PC2 value, making it one of the key residues in the formation of this domain. The second identified domain is simply comprised of the polyQ N-terminal SLiM, denoted by blue in the illustration. As the temperature approaches 415K, the $H_{aro}$ region is spatially associated with a patch of residues near the N terminus, while the domain encompassing the polyQ-adjacent SLiM expands to include all three SLiMs closest to the polyQ, including the SLiM which associated with $H_{aro}$ at 290K. For the 0Q system, PC2 (Fig. S1D) again contains two distinct domains, one around $H_{aro}$ and another around the polyQ-adjacent SLiMs. At 415K, PC2 of 0Q virtually matches that of 7Q, signaling converging dynamics for PC2 just as was observed for PC1. Based on the contact analysis performed on these ELF3-PrD ensembles, we isolate regions of interest including $H_{aro}$, the three short regions of helical propensity N-terminal to the variable polyQ domain, and F527 and proximal residues.

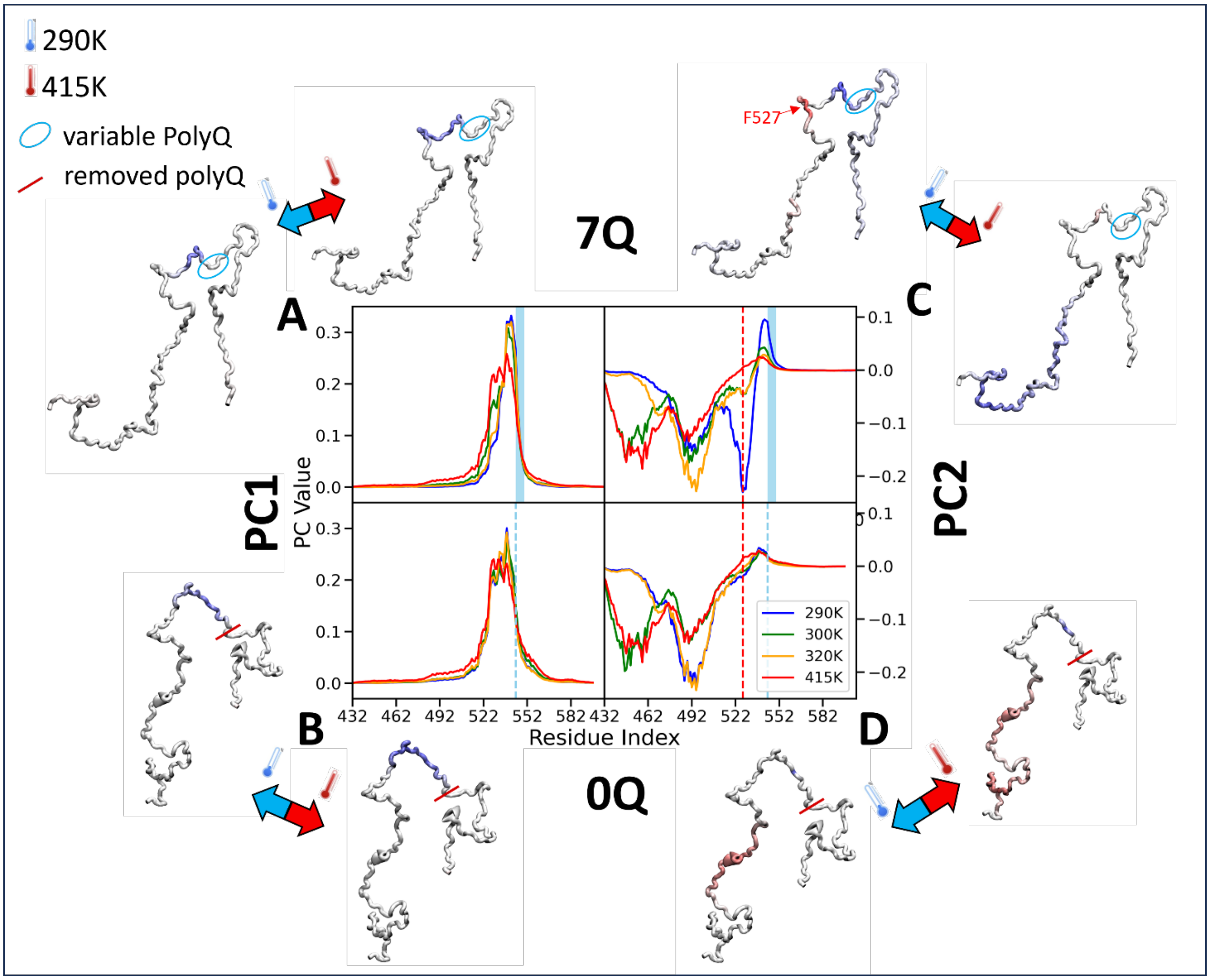


**Figure S1. A.** I-PC1 values of 7Q are plotted for each temperature in the bottom-left panel. Below this is an illustration of the ELF3-PrD with I-PC1 values mapped to each residue. 290K is on the left and 415K is on the right. PolyQ is circled in blue. **B.** I-PC1 values of 0Q are plotted for each temperature in the top-left panel. Above is an illustration of the ELF3-PrD with I-PC1 values mapped to each residue. **C.** I-PC2 values of 7Q are plotted for each temperature in the bottom-right panel. Below is an illustration of the ELF3-PrD with I-PC2 values mapped to each residue. **D.** I-PC2 values of 0Q are plotted for each temperature in the bottom-right panel. Above is an illustration of the ELF3-PrD with I-PC2 values mapped to each residue.

**Spatial Segregation of WT 7Q and 0Q REST2 Simulations as Revealed by I-PCA**

In order to understand the effect of the polyQ tract on the spatial compartmentalization of ELF3-Prd, we performed I-PCA on our 0Q and 7Q trajectories to identify spatially associated regions. Fig. S2 shows 3D representations of each system with PC values mapped onto the alpha carbons of each residue. This data is plotted in 2D in Fig. S3. At 290K, the WT PC1 (Fig. S2A) supports our previous finding that interactions between $H_{aro}$ and F527 are key to the dynamics at lower temperatures. Residues of $H_{aro}$, F527 and Y526 are shown in blue (positive PC value) indicating a propensity for interaction. Fig. S2B illustrates PC2 of the WT in which $H_{aro}$ forms a domain with the residues immediately preceding it rather than F527, and a second domain is comprised of the two SLiMs N-terminal to polyQ and some residues N-terminal to the third SLiM.

At higher temperatures, the contact dynamics of the WT begin to mirror those of the 0Q system, which is relatively temperature-invariant. Interactions of aromatic residues dominate PC1 of 0Q at both 290K and 405K (Fig. S2 D and E) with positive values being mostly aromatic with some methionine and proline residues and the trend slightly intensified at 405K. At 405K, WT mimics 0Q, with the primary driver of protein contacts coming from aromatic residues, a result corroborated by SASA results reported in a previous section. PC2 of the 0Q system highlights the 0Q-native helix (residues 554 to 566) and $H_{aro}$ + preceding residues as two independent domains with the 0Q-native helix signal weakening with temperature, corresponding to the disruption of this helix at higher temperatures. These results support the conclusion that the contact dynamics of ELF3-PrD with variable polyQ converge to that of 0Q at high temperature.

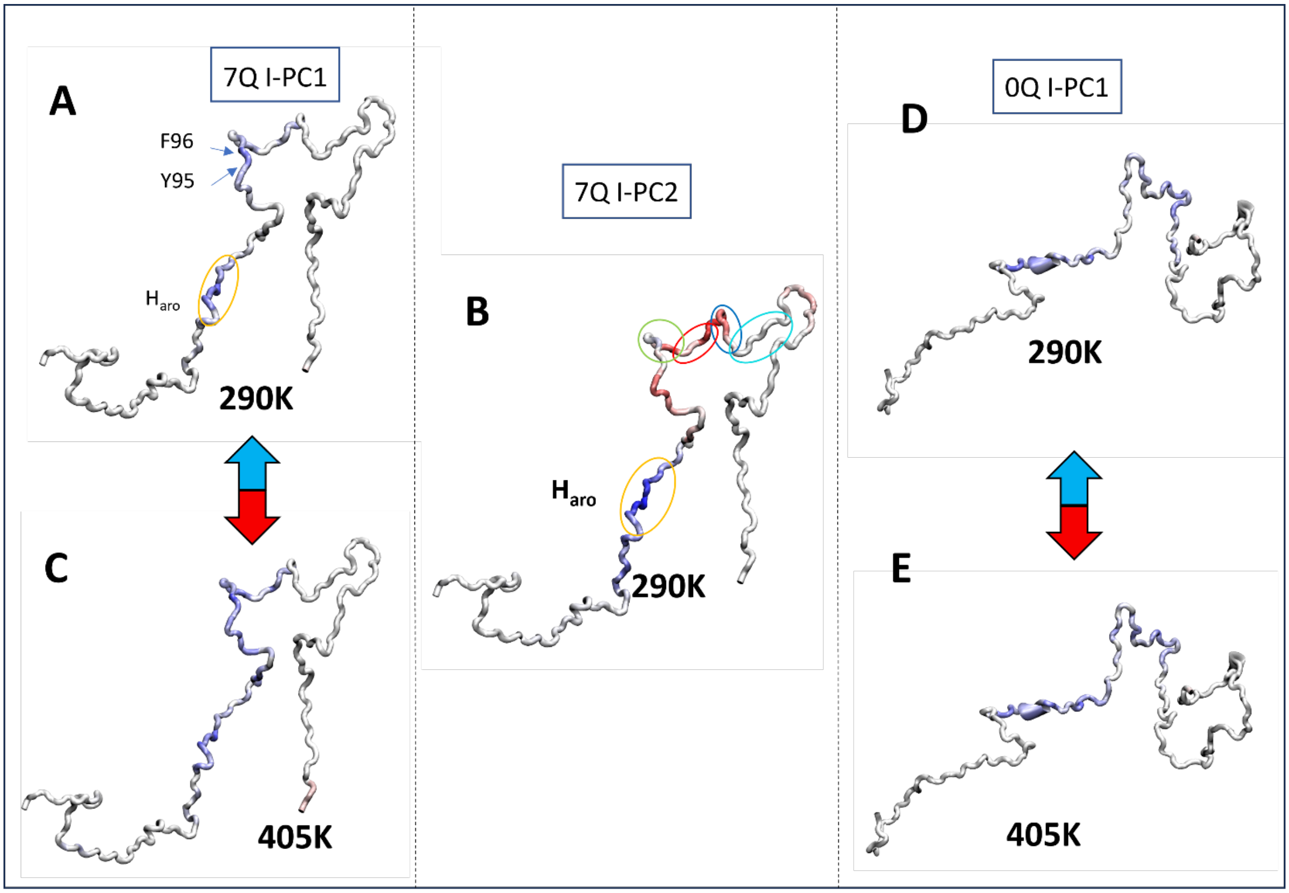


**Figure S2.** I-PCA results from REST2 trajectories are projected onto ELF3-PrD structures. **A.** I-PC1 values for the REST2 trajectory of the 7Q system at 290K are projected onto a structure of the ELF3-PrD. **B.** Projection of I-PC2 values from 7Q at 290K. **C.** Projection of I-PC1 values for 7Q at 405K. **D.** I-PC1 values from the 0Q system at 290K are projected onto a PrD structure. **E.** I-PC1 projections for 0Q at 405K.

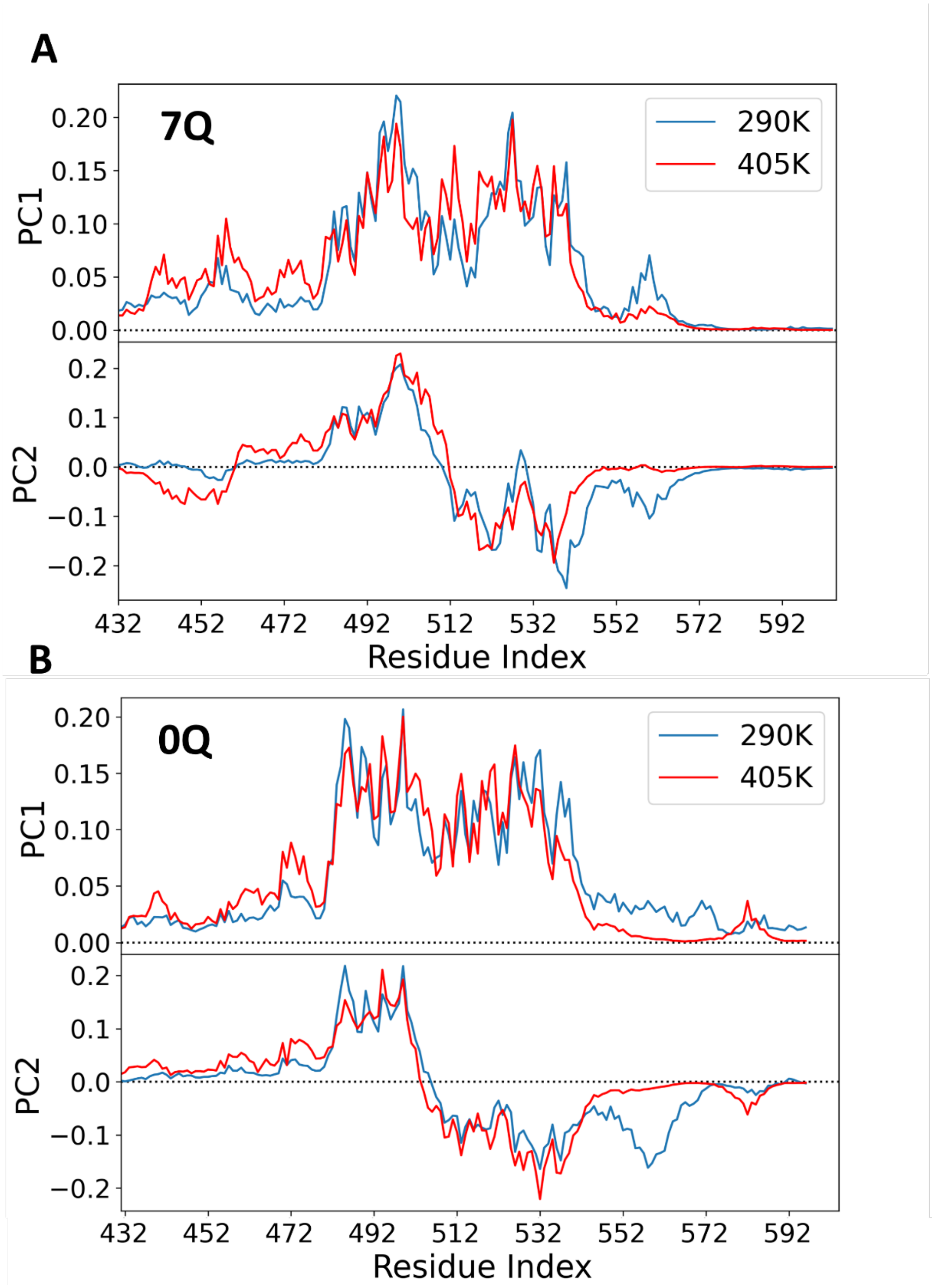


**Figure S3. A.** I-PC1 and I-PC2 values are plotted against residue index for the 7Q REST2 trajectory at 290K and 405K. **B.** I-PC1 and I-PC2 values for the 0Q REST2 trajectory at 290K and 405K.

#### Further characterization and visualization of ELF3-PrD REST2 ensembles

##### Visualization of ELF3-PrD conformational landscapes with ELViM

The Energy Landscape Visualization Method (ELViM) is a multidimensional projection technique developed to generate intuitive representations of the conformational landscapes of biomolecules. It uses intra-protein residue-residue distances as a measure of similarity between conformations. This is especially useful in the case of intrinsically disordered proteins, because, unlike some methods, no reaction coordinate needs to be defined, and unlike natively folded proteins, IDPs often lack an obvious descriptive reaction coordinate. Distances between conformations are used to create a dissimilarity matrix which is then used to project an effective two-dimensional conformational phase space. In this projection, each dot represents a conformation, and the pairwise distance between dots optimally reflects their structural similarity. A more complete description of this method is described here[4].

##### Comparing wild-type and mutant conformations in a common space

Using the ELViM method allows us to visualize the conformational landscape of the protein in a way that is particularly useful for IDPs, where specific conformations matter less than the coverage and frequency of states. Here, four temperatures of both the wild-type and F527A mutant are projected onto a common space. Figure S10A illustrates this space for both the WT and F527A mutants, where, the color of each dot is determined by the radius of gyration of the conformers. Temperature-driven trends in the conformational space can be spotted in the wild-type by observing which regions become more or less dense as the temperature increases (Figure S10B and S10C). By referencing the coloration in Fig. S4A we see that at higher temperatures, there is a slight shift towards more extended regions in the WT PrD, which can be observed as an enhancement of conformations in the upper middle region where green, yellow and red dots indicate high RG values (Fig S4B). The F527A mutant (Figure S4C) populates much different regions of conformational space than the wild-type, and the ELViM method captures this difference well. While there is some overlap, these two ensembles largely occupy different spaces, which is a testament to the drastic impact of F527 on the structures explored by ELF3-PrD. Unsurprisingly, many of the conformers explored by the F527A mutant are found in the high RG space with fewer points observed in the blue region than for the green, yellow or red regions.

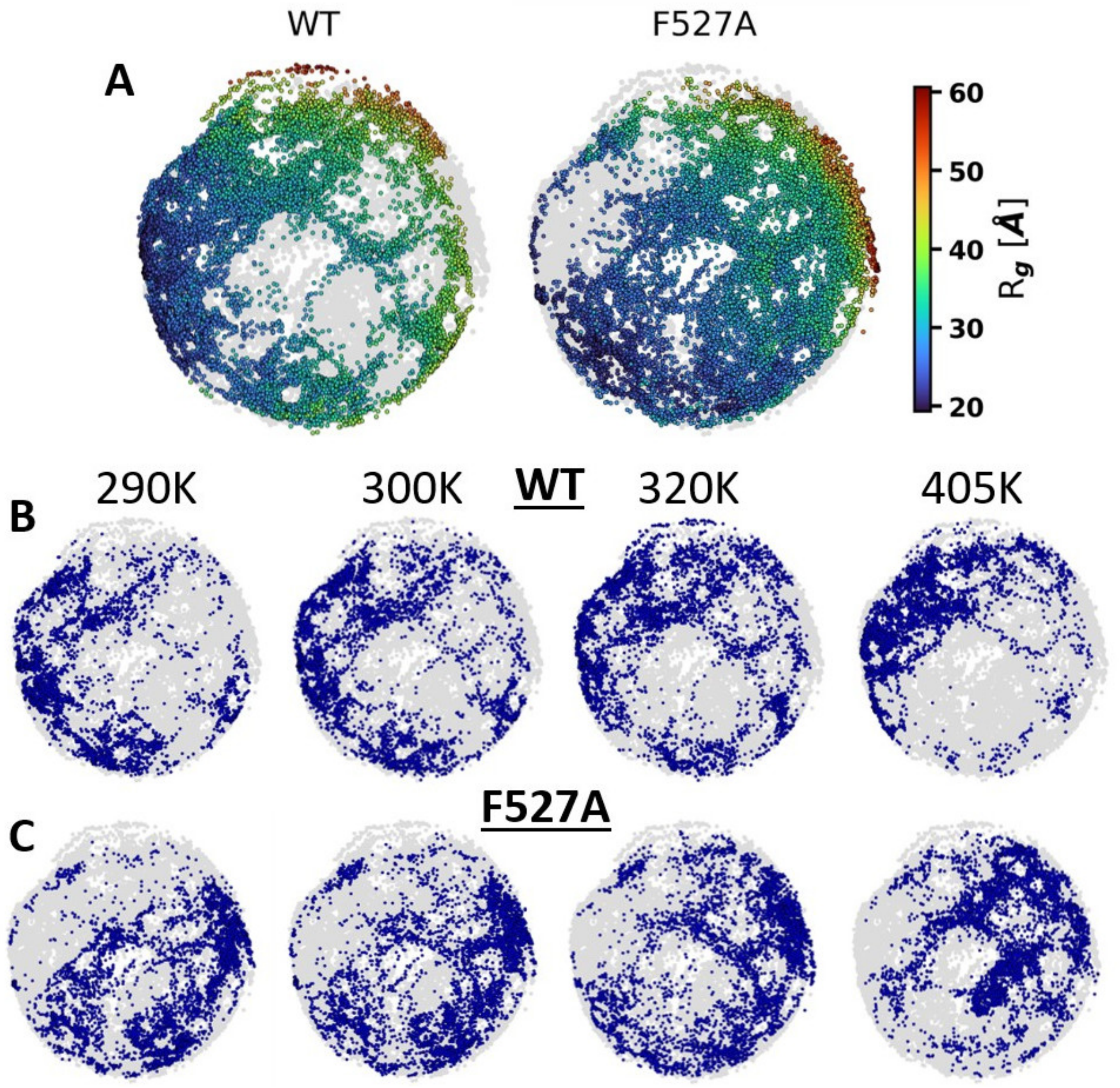


**Figure S4. A.** All conformations from WT (left) and F527A (right) systems from replicates representing approximate temperatures of 290K, 300K, 320K and 405K are mapped onto a common space using the ELViM method with each conformer colored according to radius of gyration. **B.** Population of states of wild-type ensembles with temperature increasing from left to right. **C.** Population of states for F527A with temperature increasing from left to right.

**$H_{aro}$ helicity and F527 interaction**

Once

the ELViM projection is obtained, any collective variable (CV) can be used to color the dots in the projection, revealing how CV values evolve throughout the conformational space. An example of this approach is shown in Fig. S5, where the conformational space is colored using the helicity of $H_{aro}$. The WT system has significantly more orange and red points than F527A, which is mostly comprised of blue and green points indicating that the WT conformers are more likely to exhibit helices in the $H_{aro}$ region. This could suggest that F527 does not just interact with $H_{aro}$ frequently, but that it stabilizes the secondary structure in this region.

We hypothesized that the contact status of $H_{aro}$ and F527 is important for temperature sensing because it modulates the conformational exploration of ELF3-PrD and promotes cluster formation in a thermo-sensitive manner. Therefore, we chose The $H_{aro}$/F527 minimum distance as a collective variable to visualize the prevalence of this contact in the WT and the extent to which it is disrupted in F527A. Figure S6 illustrates not only how F527A changes the sampling of the landscape, but also the extent to which they are differentiated by the interaction between F527 and $H_{aro}$. Most points in the WT space are dark blue, indicating less than 10Å distance, while the majority of points in the F527A ensemble indicate a clearly broken contact.

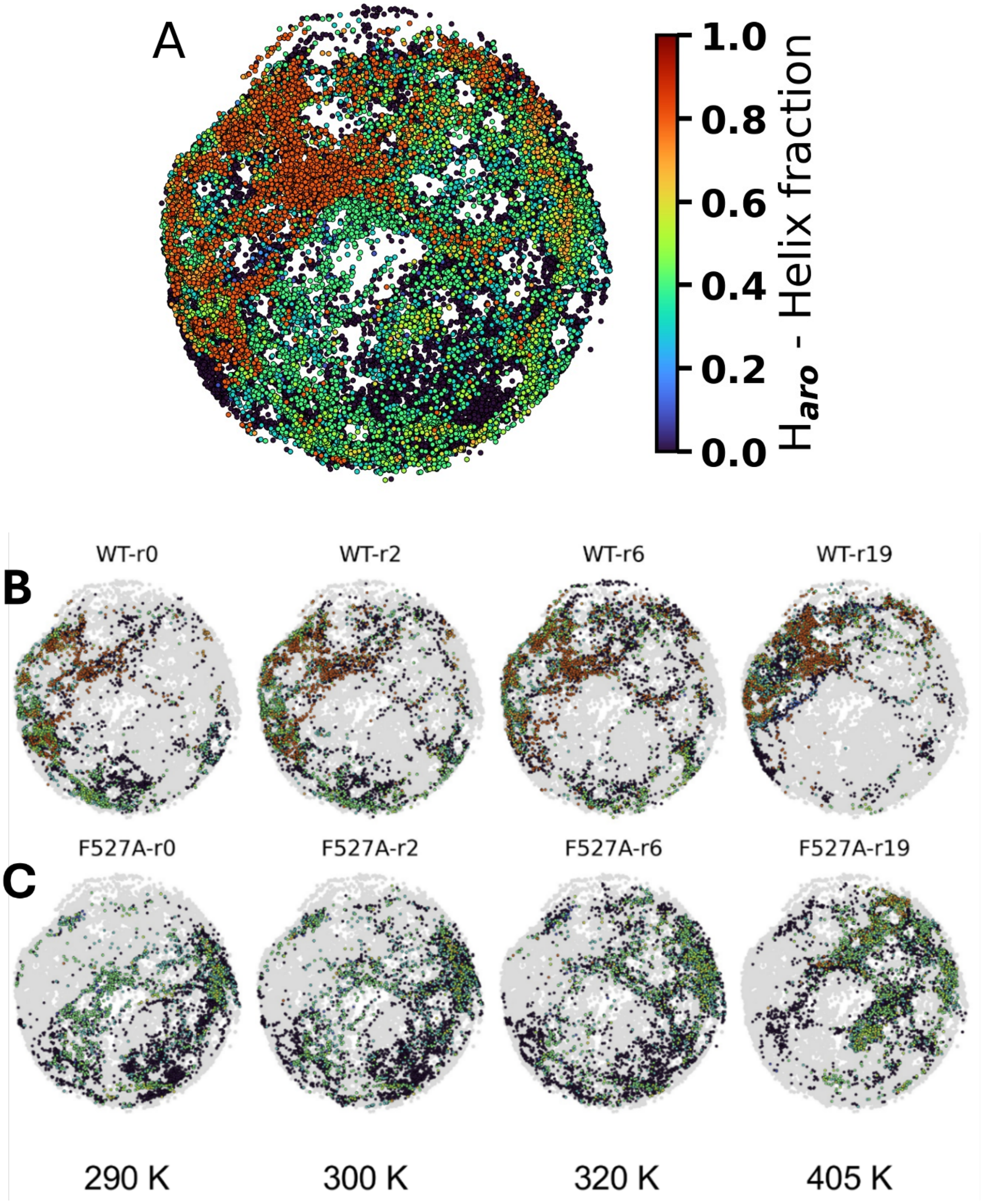


**Figure S5. Top row.** ELViM plot of the conformation space explored by WT and F527A trajectories over four approximate temperatures: 290K, 300K, 320K, and 405K. The color of the points indicates the degree of helicity in the $H_{aro}$ motif. **Center row.** Conformers from the wild-type simulations increasing in temperature from left to right. **Bottom row.** ELViM representation of F527A mutant conformations.

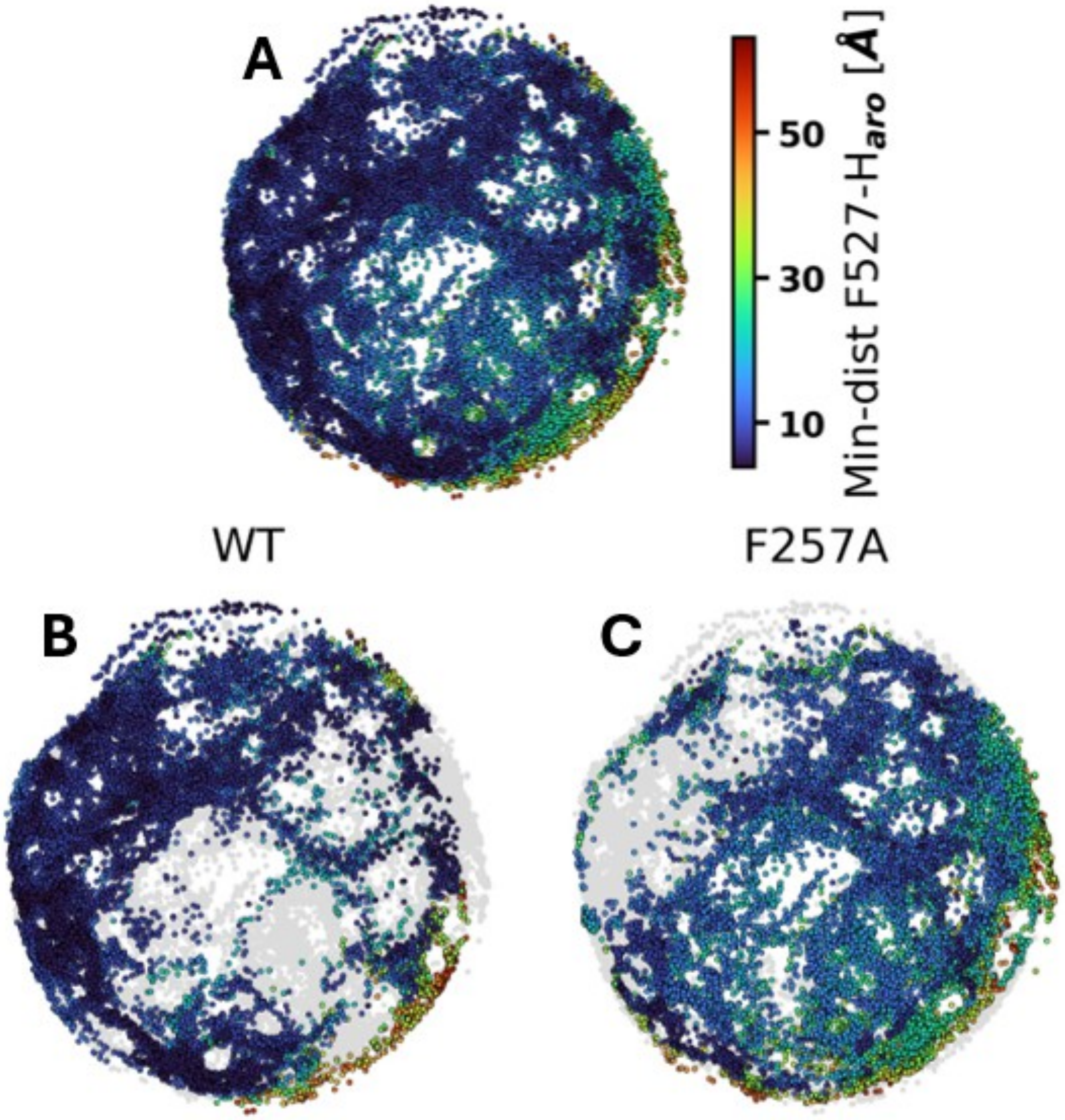


**Figure S6. A.** ELF3-PrD conformers from WT and F527A systems projected onto an ELViM space using the minimum distance between $H_{aro}$ and residue 527 as the collective variable. **B.** Conformers from 290K, 300K, 320K and 405K simulations of WT projected independently on the ELViM space, again using the $H_{aro}$/residue 527 minimum distance as a collective variable. **C.** The F527A conformational space projected in the same way as the WT.

### Supplementary Figures

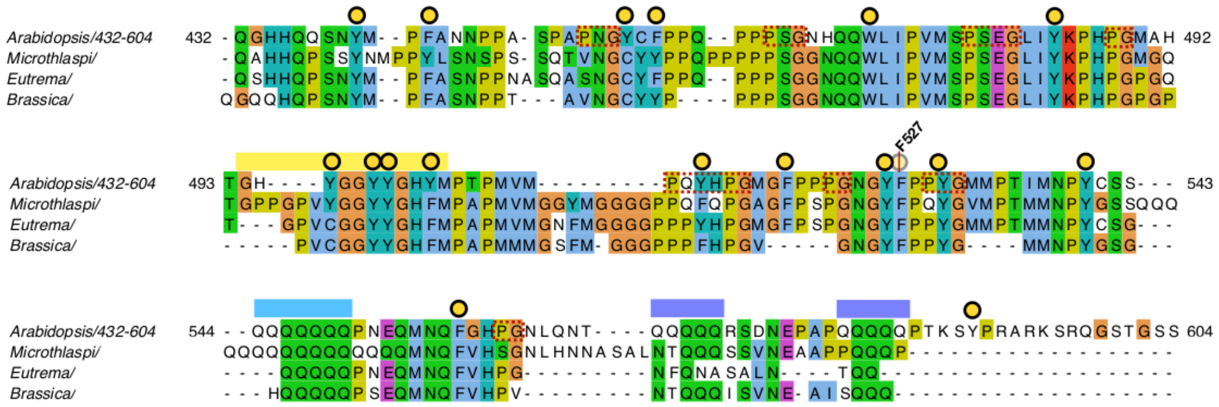


**Figure S7.** A sequence alignment of the prion-like domain of ELF3 is shown. Yellow dots indicate aromatic residues. Pro-$X_n$-Gly motifs are denoted by dashed red boxes.

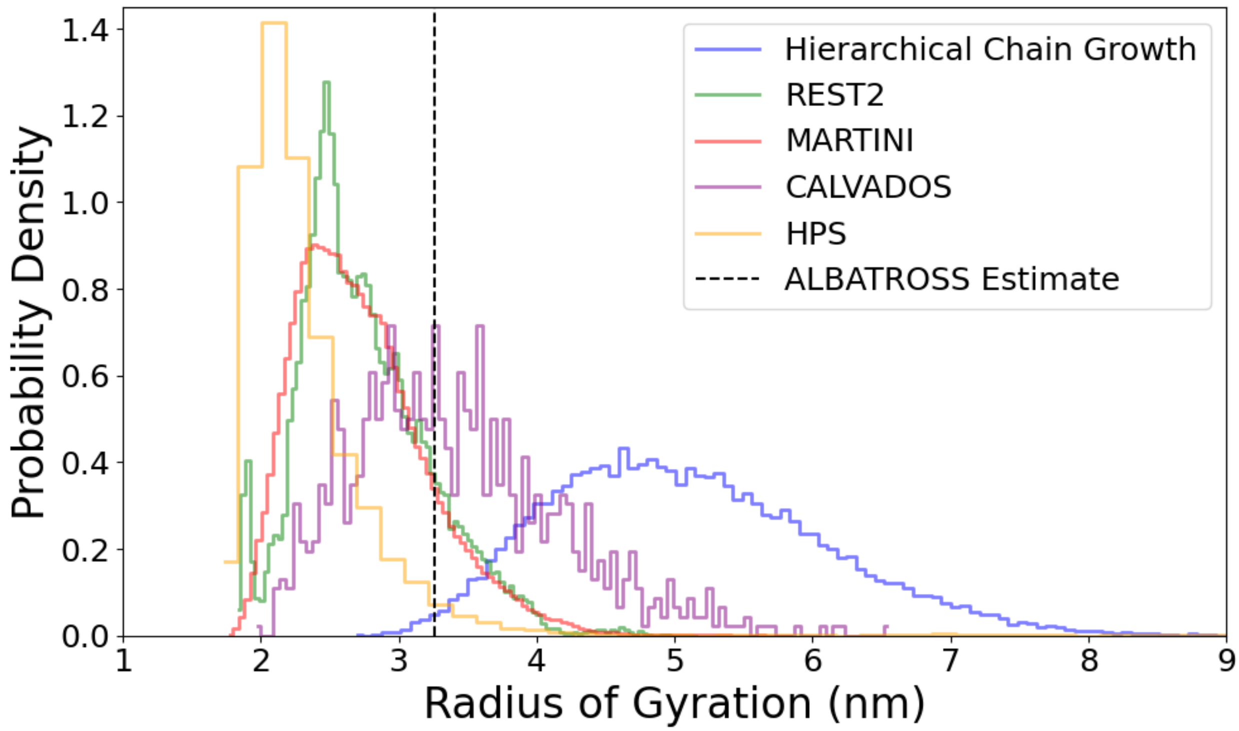


**Figure S8.** Radius of gyration distributions obtained by six different methods are shown. The black dashed line indicates the average RG value calculated by ALBATROSS, which does not compute RG as an ensemble property.

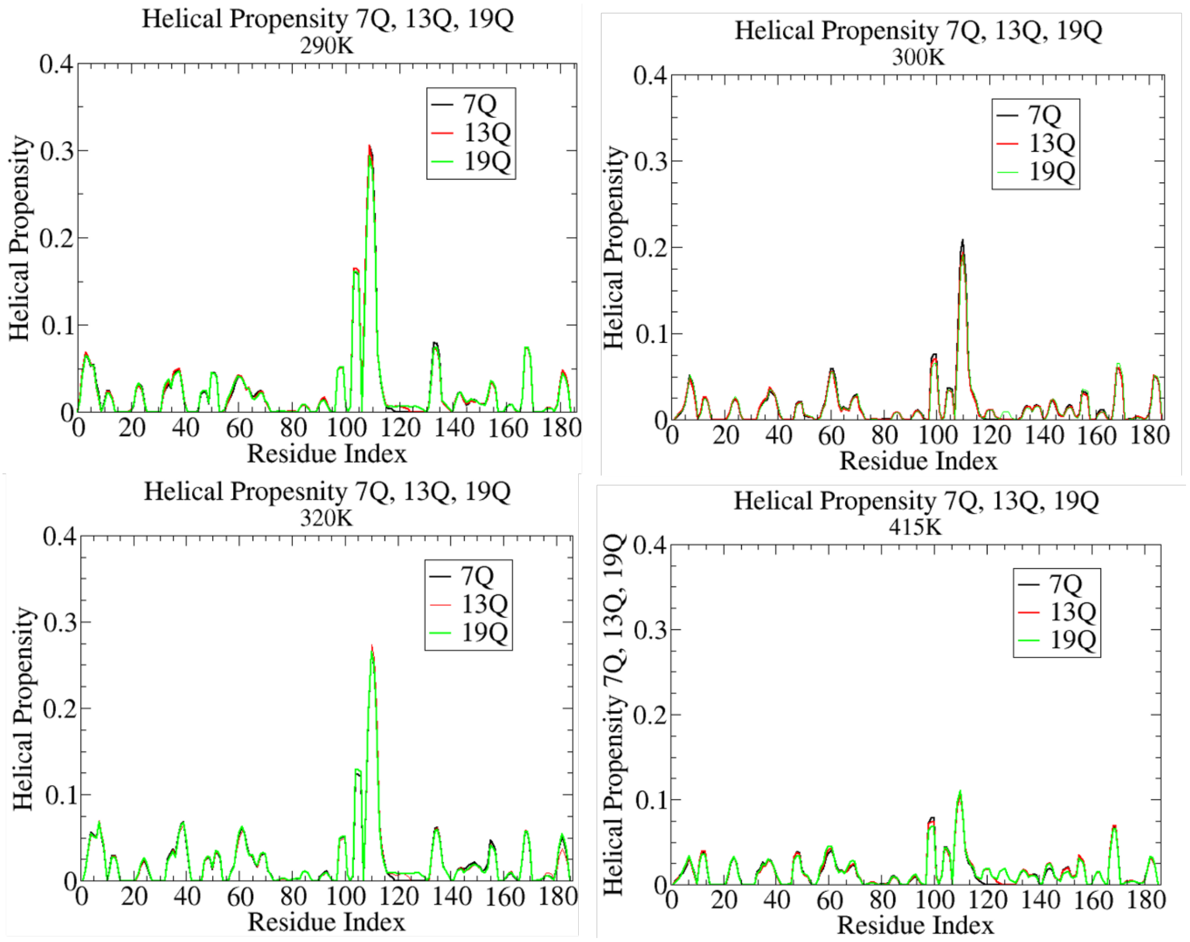


**Figure S9.** Helical propensity of HCG ensembles at four temperatures.

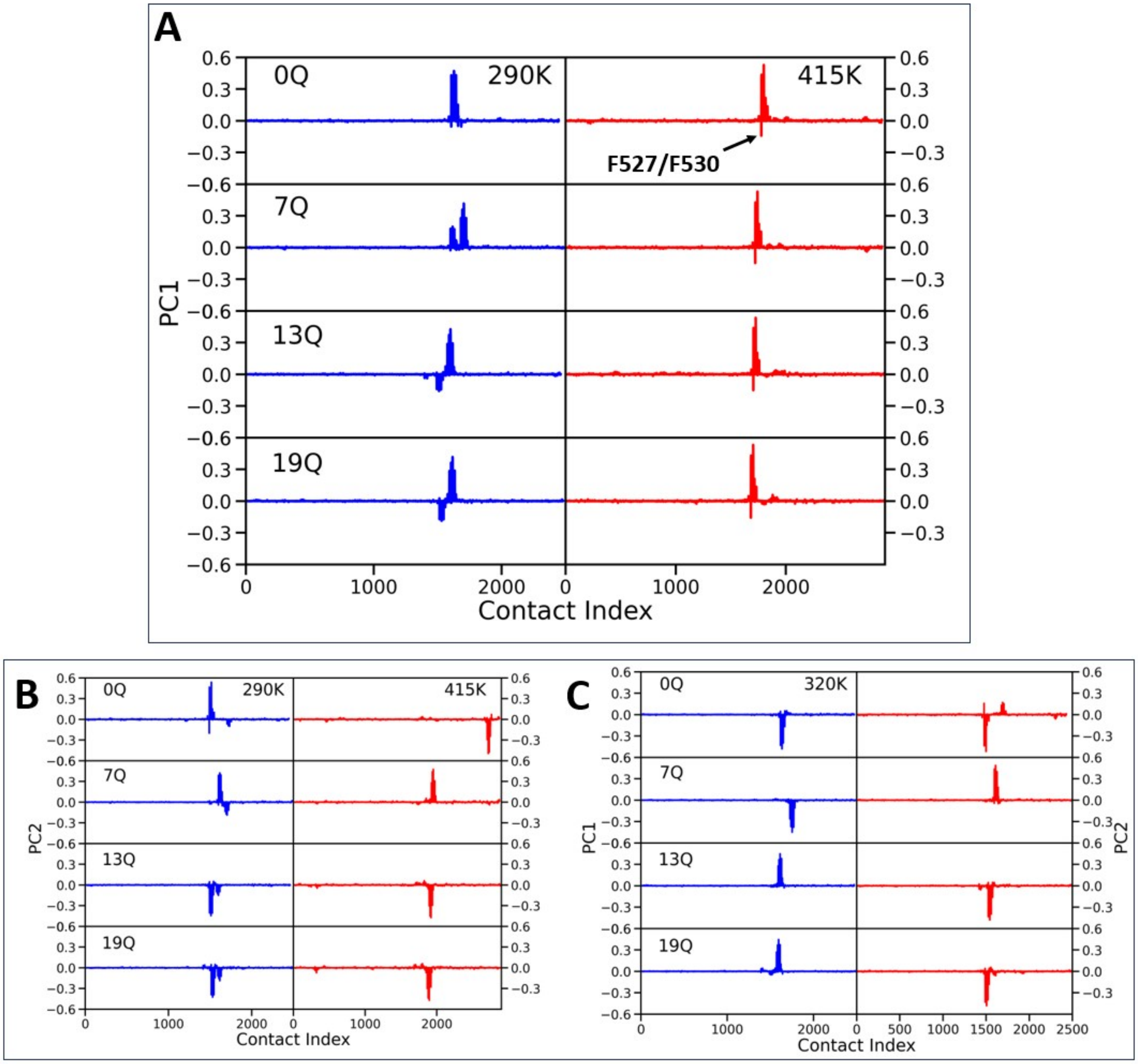


**Figure S10. A.** The top PC from E-PCA of HCG ensembles at 290K (left) and 415K (right). **B.** The second PC from E-PCA of HCG ensembles at 290K (left) and 415K (right). **C.** The top two PCs from E-PCA of the HCG ensembles at 320K.

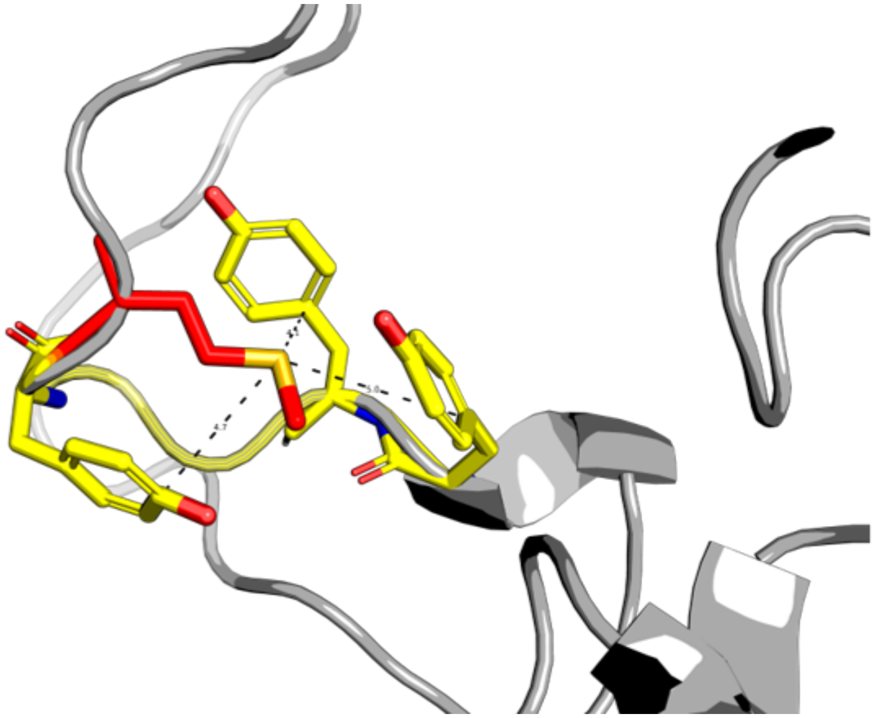

**Figure S11.** Methionine 504 coordinating interactions with three tyrosine residues.

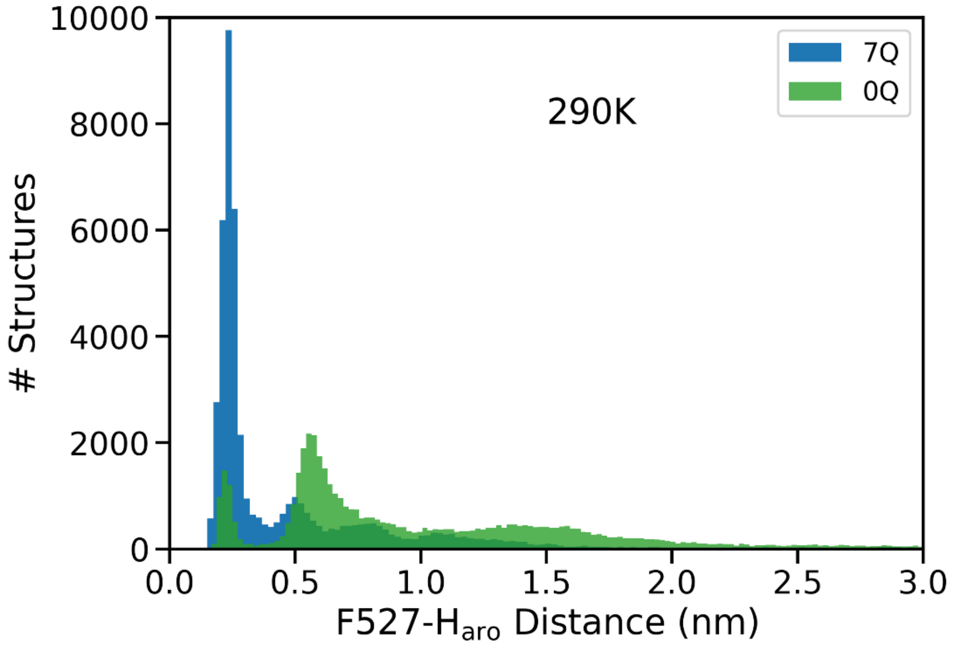


**Figure S12.** Distribution of distance between $H_{aro}$ and F527 for the 7Q and 0Q systems.

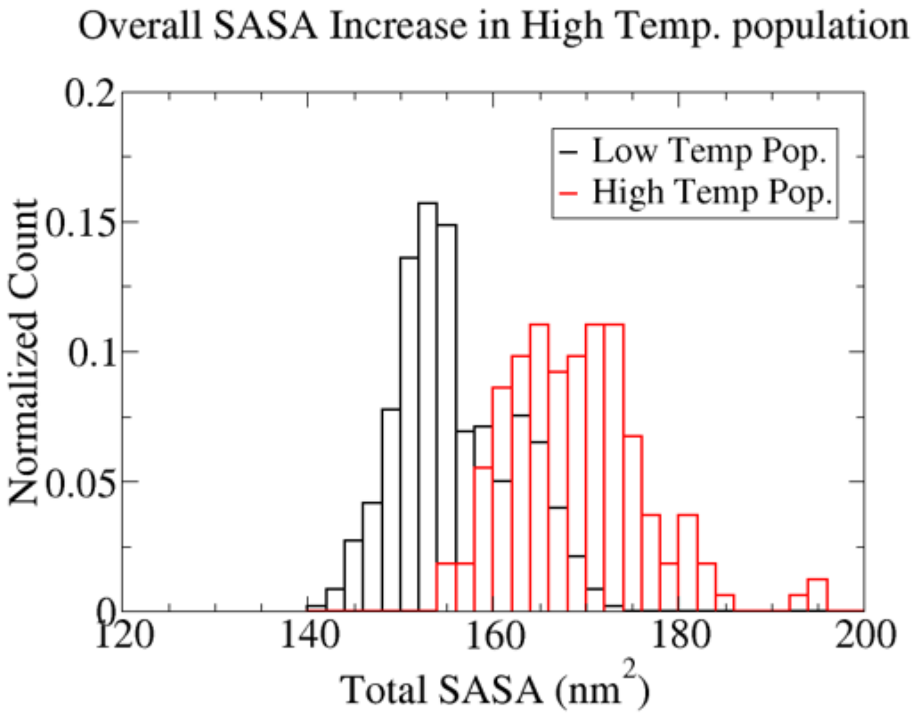


**Figure S13.** Total solvent accessible surface area for the low and high temperature populations as identified in Fig. 3.

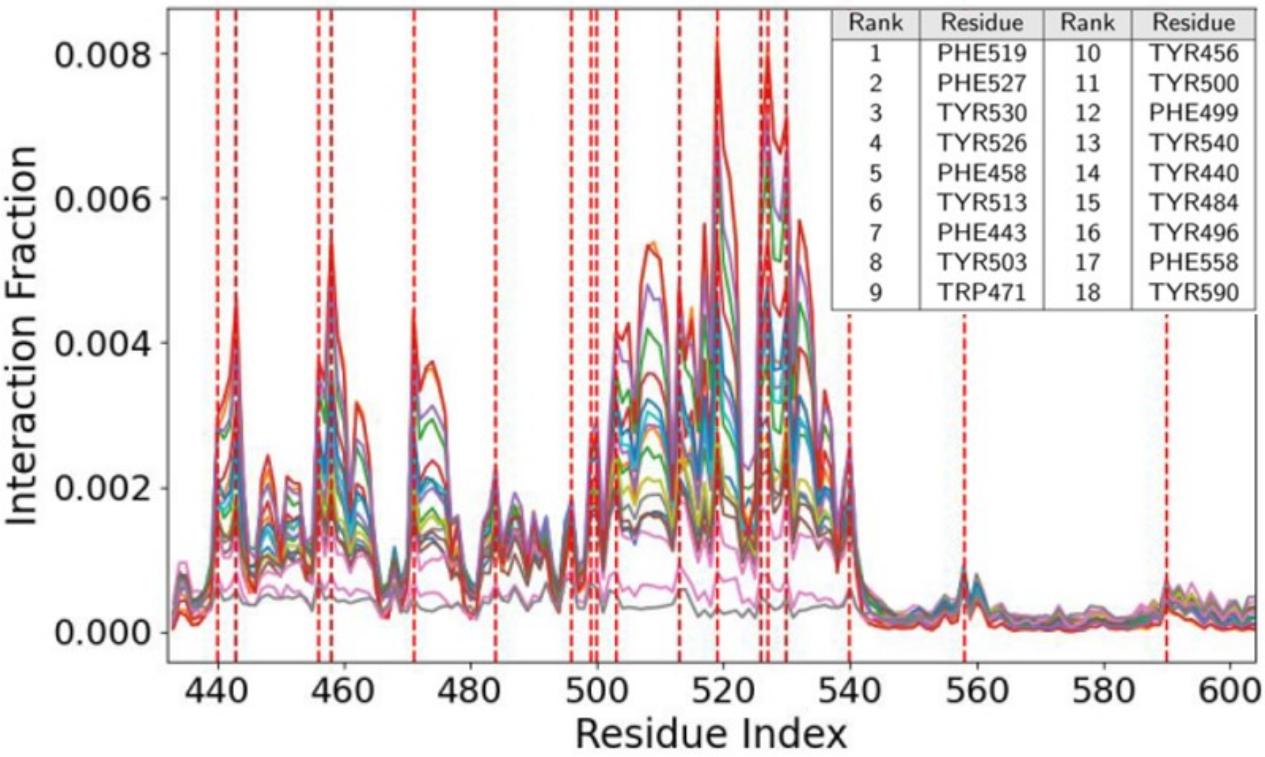


**Figure S14. Inter-protein interaction profiles of all 18 aromatic residues of ELF3-PrD. The inset is a table ranking each aromatic by interaction frequency. Aromatic residues are indicated by red dashed lines.**

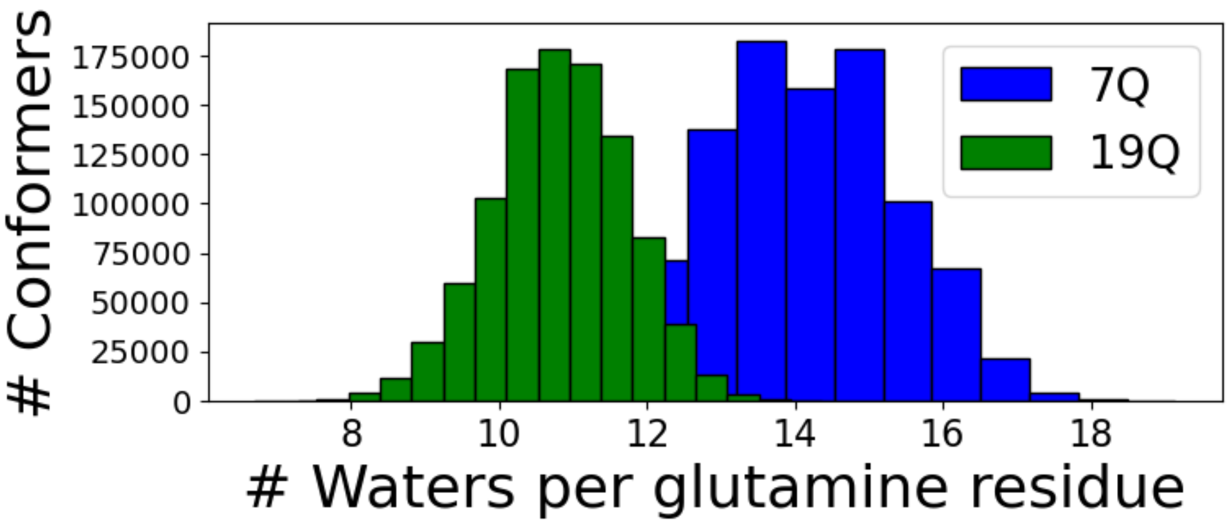


**Fig. S16.** The number of water molecules interacting with the polyQ tract normalized by number of glutamine residues for the 7Q and 19Q systems.

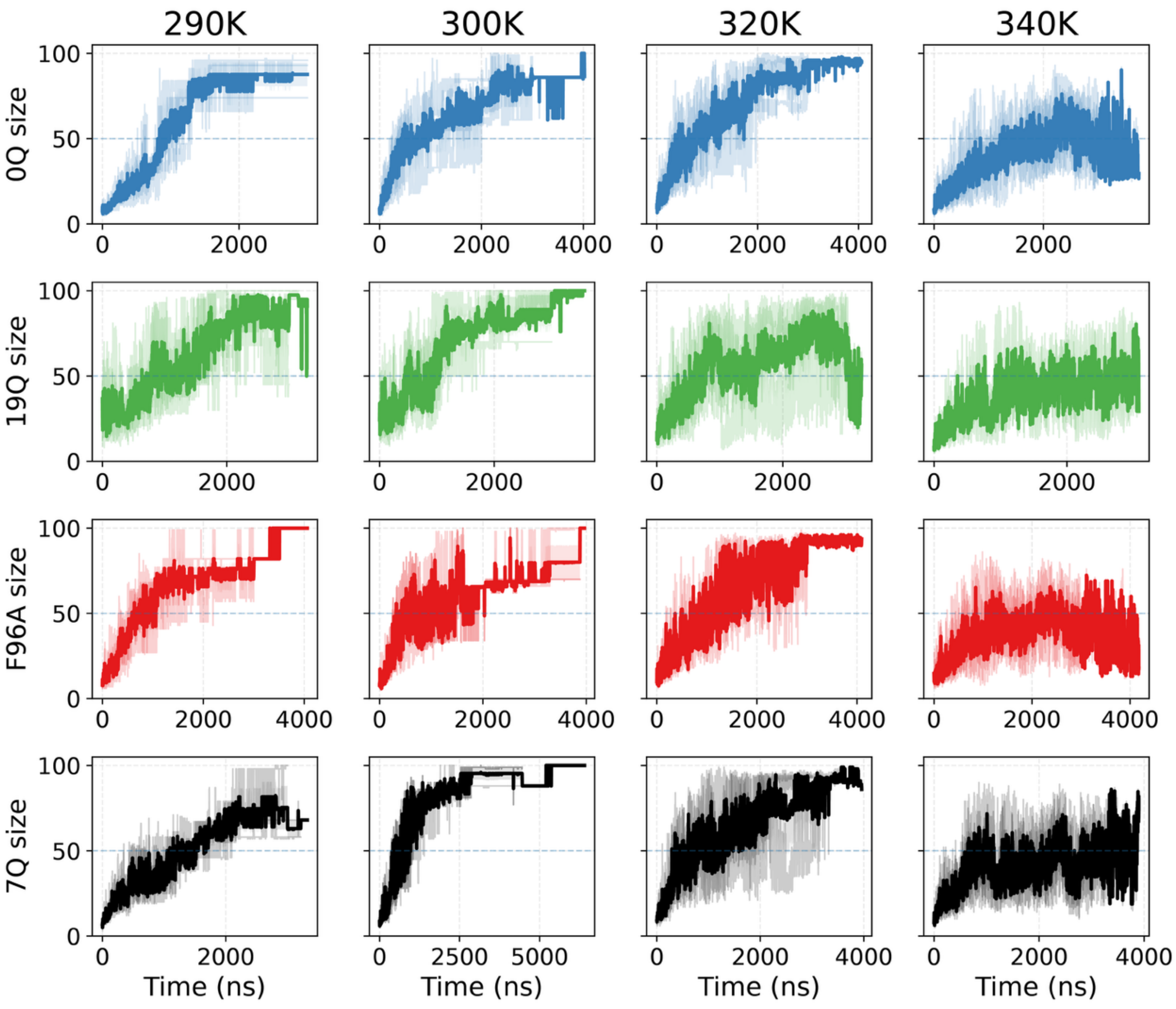


**Figure S17. Largest-cluster growth and persistence across temperature and ELF3-PrD variants.** Time series of the size of the dominant cluster, $N_{\mathrm{max}}(t)$(number of peptides in the largest cluster), shown for four variants (rows: 0Q, 19Q, F527A—labeled F96A in this plot—and 7Q/WT) across four temperatures (columns: 290 K, 300 K, 320 K, 340 K). Semi-transparent traces show three individual replicas, and the darker trace indicates the replicate-averaged trajectory. Clusters are identified using a 0.5 nm cutoff between backbone beads. The dashed horizontal line at $N_{\mathrm{max}} = 50$marks the threshold used to define a condensed state in lifetime/persistence analyses..

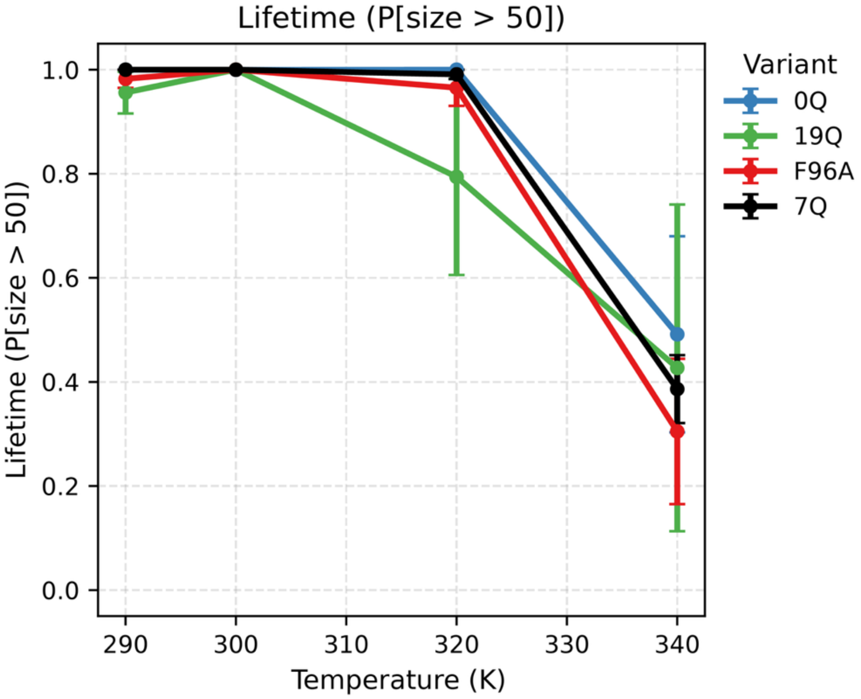


**Figure S18. Condensed-state persistence (lifetime) as a function of temperature.** Condensate lifetime defined as the fraction of simulation frames satisfying $N_{\max}(t) > 50$ (i.e., the dominant cluster contains >50 peptides), plotted versus temperature for 7Q (WT), 0Q, 19Q, and F527A. Points show replicate-averaged values and error bars indicate SEM across three independent replicas. These metric reports whether the system remains persistently condensed versus intermittently dissolving/re-forming at higher temperature.

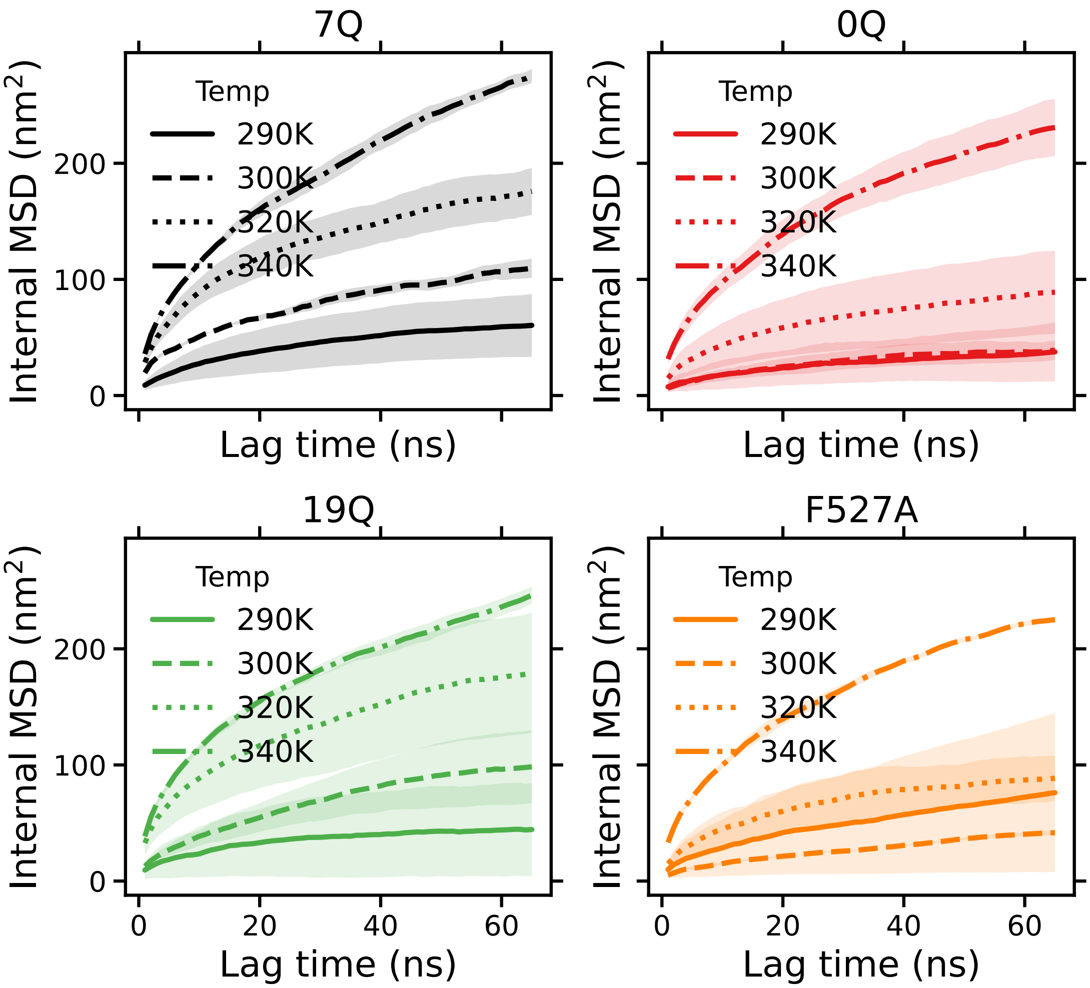


**Figure S19. Internal mobility inside the condensed phase quantified by internal MSD.** Internal mean-squared displacement (MSD; nm$^2$) versus lag time (ns) for peptides in the condensed phase, shown separately for each variant (7Q/WT, 0Q, 19Q, F527A) and temperature (290 K, 300 K, 320 K, 340 K; line styles as indicated). "Internal" MSD is computed after removing bulk motion of the condensate (i.e., reporting relative motion within the dense phase rather than center-of-mass drift). Shaded bands indicate variability across three independent replicas. Curvature/sublinear growth reflects constrained, non-Brownian internal rearrangements.

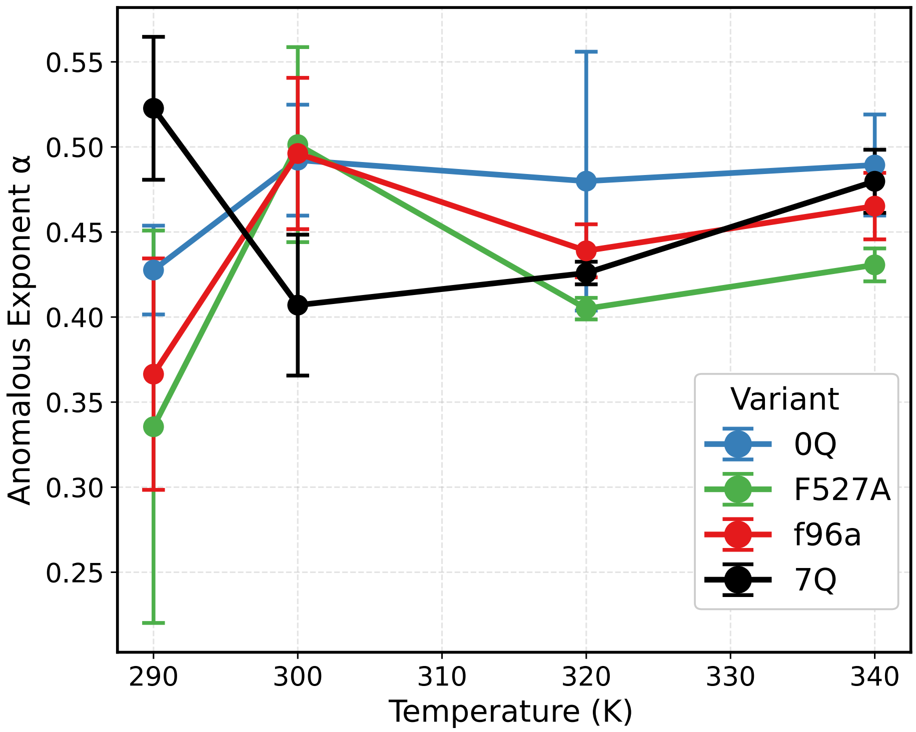


**Figure S20. Subdiffusive exponent $\alpha$ reveals viscoelastic dynamics in the condensed state.** Anomalous exponent $\alpha$ extracted from MSD scaling ($\langle \Delta r^2(\tau) \rangle \sim \tau^{\alpha}$) as a function of temperature for 7Q (WT), 0Q, 19Q, and F527A. Points show replicate means and error bars indicate SEM across three independent replicas. Values $\alpha < 1$indicate subdiffusive, dynamically constrained motion consistent with viscoelastic/heterogeneous condensate dynamics rather than an ideal Brownian liquid.

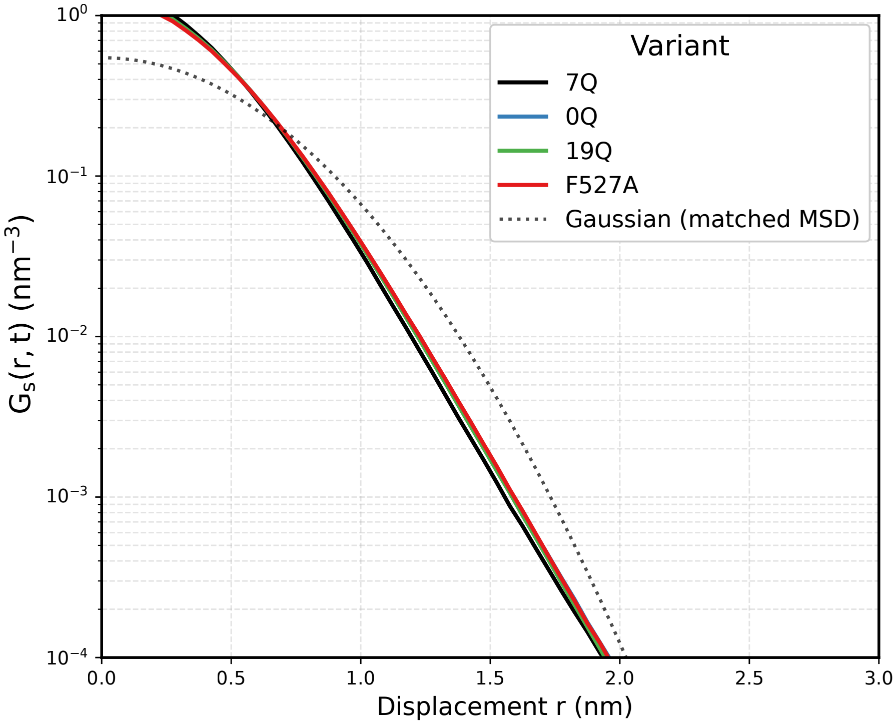


**Figure S21. Non-Gaussian displacement statistics indicate dynamic heterogeneity.** Self van Hove function $G_s(r, t^*)$at 300 K and $t^* = 1$ns for each variant (7Q/WT, 0Q, 19Q, and F527A). The dotted curve shows a Gaussian reference distribution matched to the MSD. Deviations from the Gaussian, particularly excess probability at small displacements reflect heterogeneous/caging-like dynamics within the dense phase.

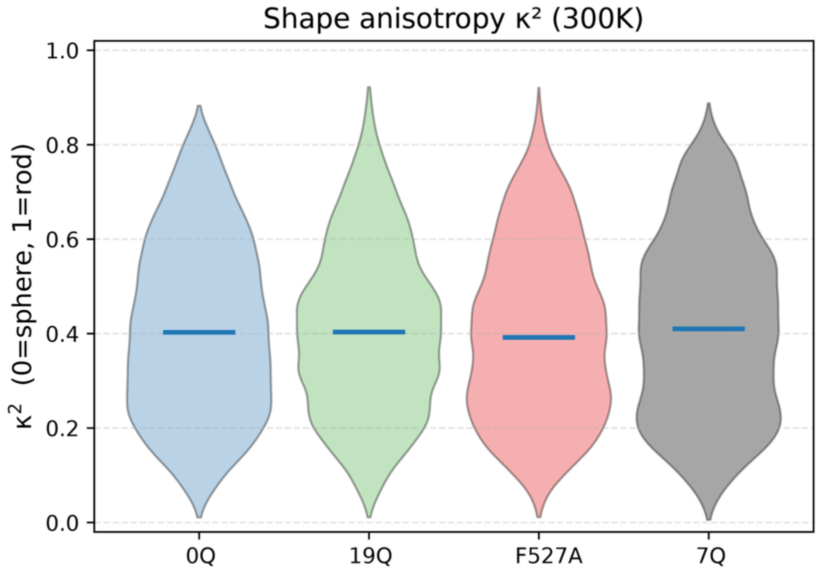


**Figure S22. Condensate morphology at 300 K quantified by shape anisotropy $\kappa^2$.** Distribution of the shape anisotropy $\kappa^2$of the dominant condensed assembly at 300 K for 0Q, 19Q, F527A, and WT (7Q). $\kappa^2 = 0$ corresponds to a spherical cluster and $\kappa^2 = 1$to a rod-like object. Violin plots show the distribution across frames (pooled across three replicas within the analysis window – last 1 micro-second of trajectory); the horizontal bar indicates the central tendency for each variant.

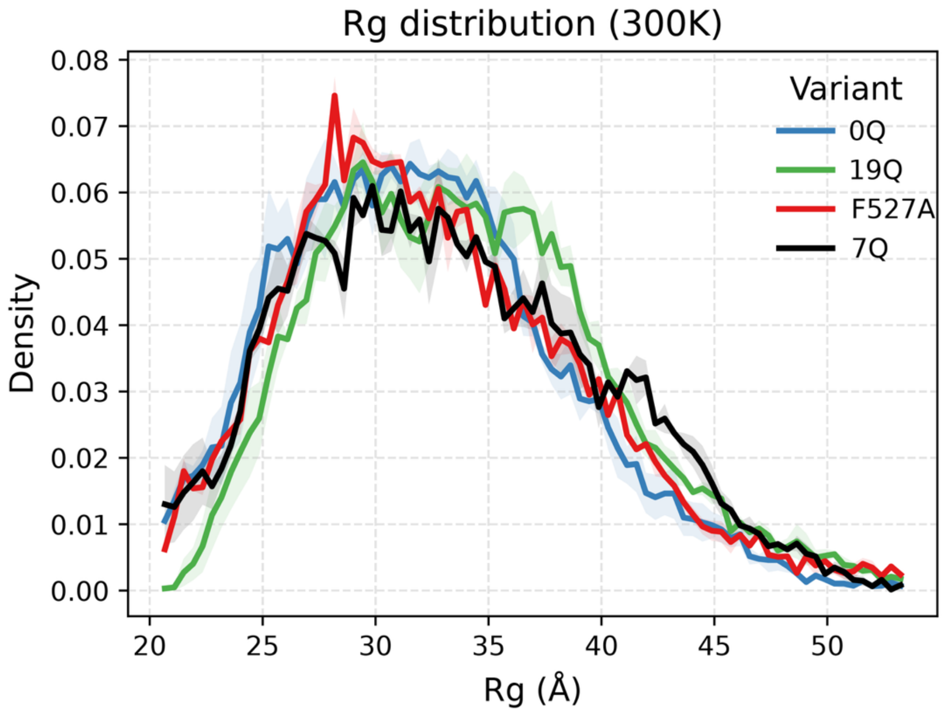


**Figure S23.** Single-chain compaction at 300 K across variants. Probability density of the per-chain radius of gyration, $R_g$(Å), at 300 K for 0Q, 19Q, F527A, and WT (7Q). Curves are computed over the analysis window – last 1 micro-second of trajectory; shaded bands indicate variability across three independent replicas. This comparison tests whether variants exhibit large differences in single-chain compaction under the same temperature conditions.

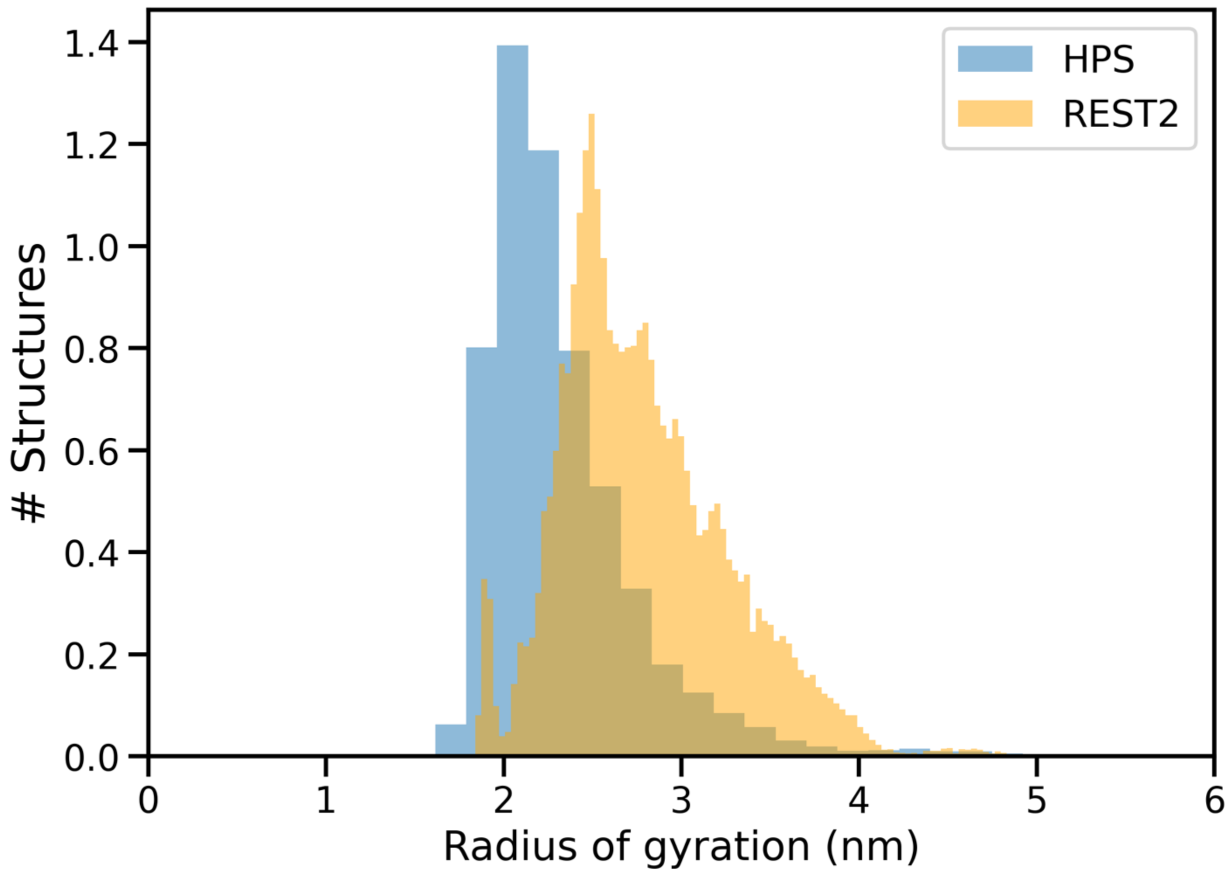


**Figure S24.** Radius of gyration comparison of ELF3-PrD between the coarse grained HPS forcefield and all-atom REST2 simulations.

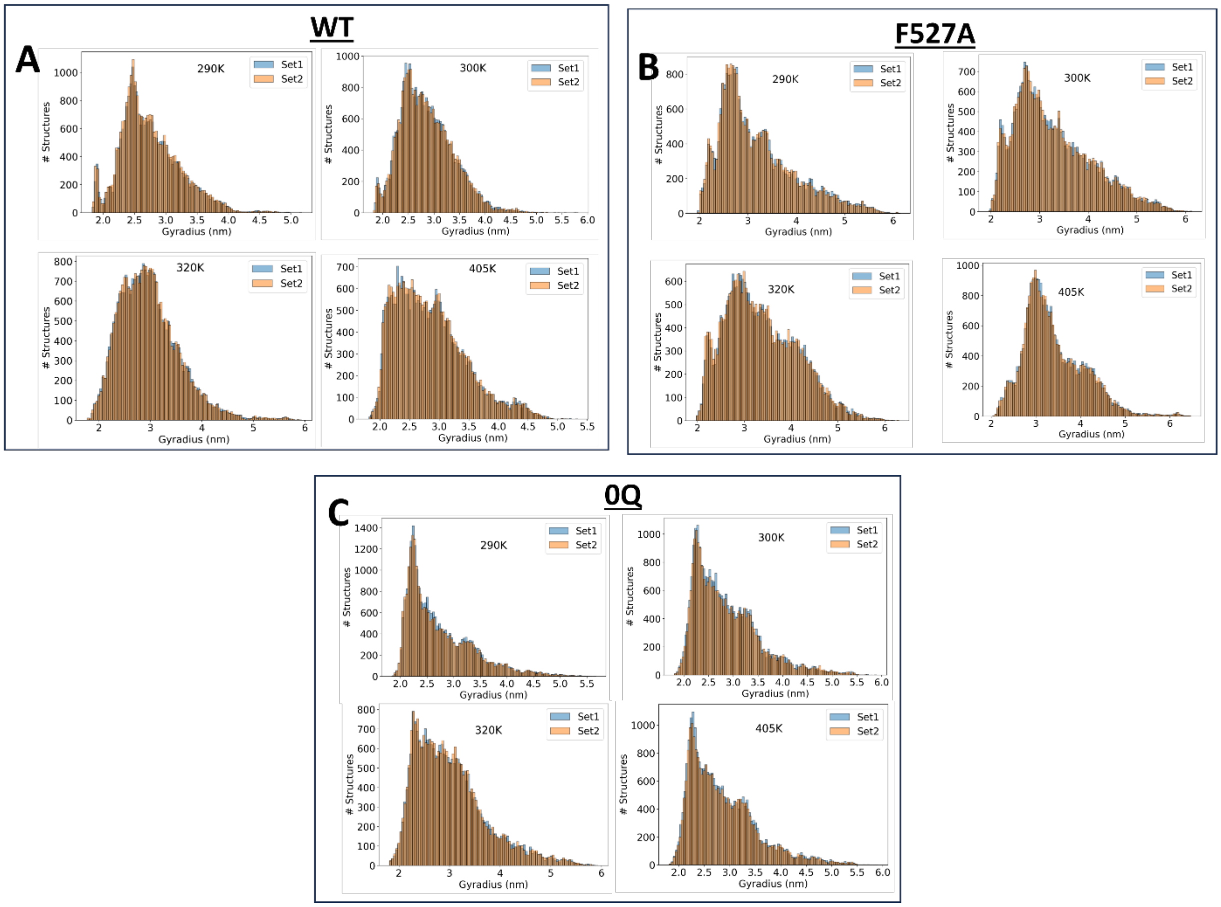


**Figure S25.** Split analyses of radius of gyration to test for convergence of REST2 simulations. **A.** Split analysis for four replicas of the wildtype trajectories where radius of gyration values from random frames have been assigned to one of two groups. **B.** Split RG analysis of four temperature replicates of the F527A mutant. Split RG analysis of four temperature replicates of the 0Q system.

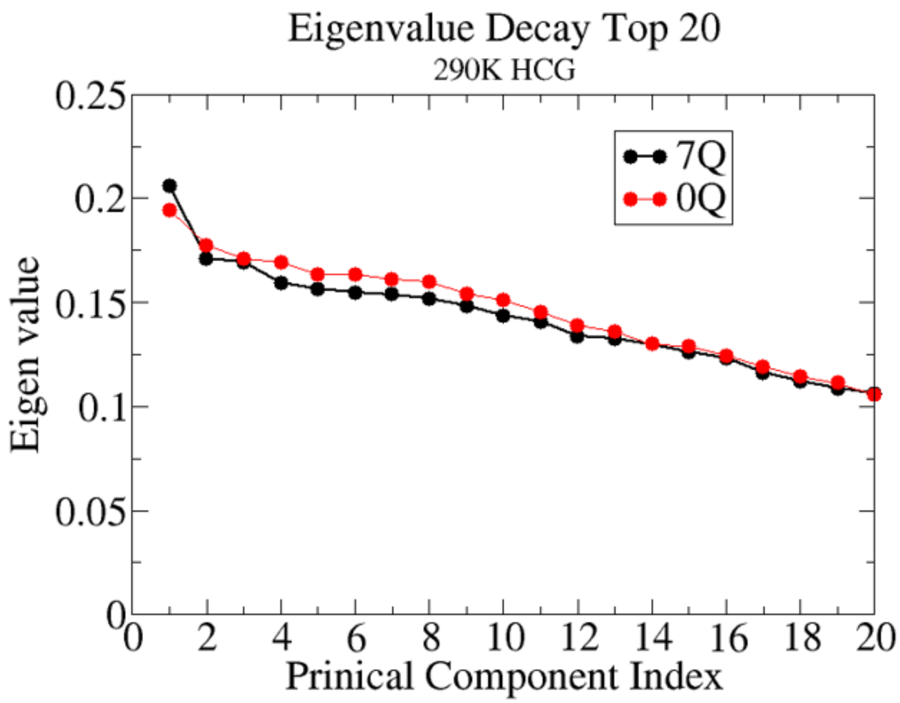


**Figure S26.** The first 20 eigenvalues obtained by performing I-PCA on 7Q and 0Q HCG ensembles at 290K.

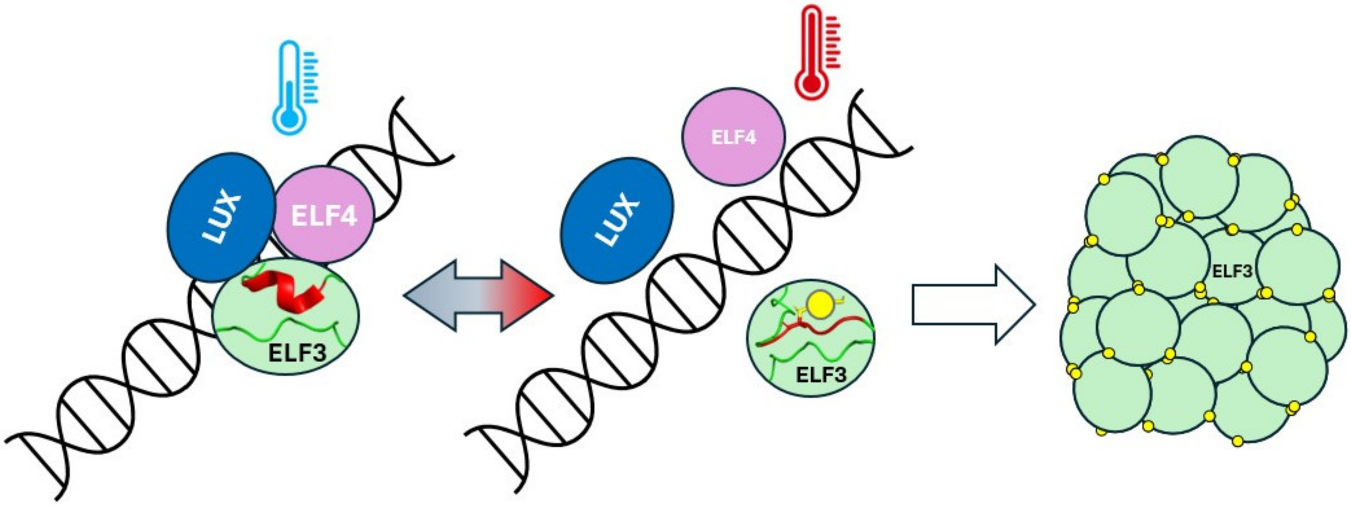


**Figure S27.** An illustration of a proposed hypothesis in which **i)** high temperature destabilizes the evening complex by melting SLiM helices, **ii)** aromatic residues are exposed and **iii)** condensate formation is promoted.

| System | RG | $R_{ee}$ | Nu | asphericity |
|---|---|---|---|---|
| WT | 3.259 | 7.5 | 0.38 | 0.329 |
| 0Q | 3.259 | 7.6 | 0.39 | 0.34 |
| F96A | 3.319 | 7.65 | 0.394 | 0.334 |
| 19Q | 3.259 | 7.47 | 0.36 | 0.31 |

**Table S1.** Predicted measurements for a variety of ELF3 PrD variants obtained from the ALBATROSS deep learning algorithm. Radius of gyration, end-to-end distance, polymer scaling exponent and asphericity are represented, respectively.

| System | Rg | $R_{ee}$ | Nu | <delta> | <S> |
|---|---|---|---|---|---|
| WT_290K | 3.4109 | 7.90 | 0.49 | 0.19 | 0.78 |
| WT_300K | 3.402 | 7.91 | 0.49 | 0.18 | 0.78 |
| WT_340K | 3.727 | 8.8 | 0.52 | 0.52 | 0.95 |
| 0Q_300K | 3.402 | 7.92 | 0.49 | 0.18 | 0.78 |
| 0Q_340K | 3.42 | 8.01 | 0.507 | 0.201 | 0.811 |
| F527A_290K | 3.313 | 7.61 | 0.47 | 0.19 | 0.75 |
| F527A_340K | 3.89 | 8.65 | 0.513 | 0.213 | 0.808 |
| 13Q_290K | 3.379 | 7.84 | 0.480 | 0.187 | 0.699 |
| 13Q_340K | 3.726 | 8.764 | 0.514 | 0.193 | 0.856 |
| 19Q_290K | 3.695 | 8.76 | 0.509 | 0.2212 | 0.831 |
| 19Q_340K | 3.798 | 8.93 | 0.513 | 0.196 | 0.84 |

**Table S2.** Predicted measurements for a variety of ELF3 PrD variants obtained from the CALVADOS2 web server. Radius of gyration, end-to-end distance, polymer scaling exponent and SAXS scattering curves.
